# Glutamatergic dysfunction precedes neuron loss in cerebral organoids with *MAPT* mutation

**DOI:** 10.1101/2021.02.03.429623

**Authors:** Kathryn R. Bowles, M. Catarina Silva, Kristen Whitney, Taylor Bertucci, Jacob C. Garza, Nathan C. Boles, Kevin H. Strang, Sidhartha Mahali, Jacob A. Marsh, Cynthia Chen, Derian A. Pugh, Yiyuan Liu, Joshua E. Berlind, Jesse D. Lai, Susan K. Goderie, Rebecca Chowdhury, Steven Lotz, Keith Lane, Khadijah Onanuga, Celeste M. Karch, Justin K. Ichida, John F. Crary, Stephen J. Haggarty, Alison M. Goate, Sally Temple

## Abstract

Frontotemporal dementia (FTD) due to *MAPT* mutation causes pathological accumulation of tau and glutamatergic cortical neuronal death by unknown mechanisms. We used human induced pluripotent stem cell (iPSC)-derived cerebral organoids expressing tau-V337M and isogenic corrected controls to discover early alterations due to the mutation that precede neurodegeneration. At 2 months, mutant organoids show upregulated expression of *MAPT*, and glutamatergic signaling pathways and regulators including the RNA-binding protein *ELAVL4*. Over the following 4 months, mutant organoids accumulate splicing changes, disruption of autophagy function and build-up of tau and P-tau S396. By 6 months, tau-V337M organoids show specific loss of glutamatergic neurons of layers affected in patients. Mutant neurons are susceptible to glutamate toxicity which was rescued pharmacologically by treatment with the PIKFYVE kinase inhibitor apilimod. Our results demonstrate a sequence of events that precede cell death, revealing molecular pathways associated with glutamate signaling as potential targets for therapeutic intervention in FTD.

## INTRODUCTION

Frontotemporal dementia (FTD) encompasses a spectrum of disorders, accounting for 5-6% of all dementia, and up to 20% of cases under age 65 (Bird et al., 2003; Olszewska et al., 2016). FTD is heritable, with up to 50% of cases having a family history and 10-20% of these show autosomal dominant inheritance due to mutations in the microtubule-associated protein tau (*MAPT*) gene (FTD-tau) (Benussi et al., 2015; Rohrer et al., 2009). Pathologic studies reveal accumulation of filamentous, hyperphosphorylated tau protein (P-tau) in cortical and hippocampal neurons and glia, and tau redistribution from axons to the somatodendritic compartment. These changes correlate with degeneration of synapses and neuronal loss in frontal and temporal cortices (Bodea et al., 2016; Clare et al., 2010). These analyses have been performed in post-mortem tissue, and thus represent later disease stages when cell loss is already advanced. The lack of disease-modifying therapies for FTD-tau underscores the urgency to identify early molecular changes and disease biomarkers that precede overt pathology and cell death, when therapeutic intervention will likely have most impact.

As familial and sporadic FTD-tau are highly similar clinically and neuropathologically (Forrest et al., 2018), there is a good rationale for studying patients with *MAPT* mutations to further understand disease processes. *MAPT* is alternatively spliced at its C-terminus, resulting in two major isoforms, 3R and 4R tau, defined by exclusion or inclusion of exon 10, respectively. To date, 111 unique *MAPT* variants have been identified: 56 are pathogenic (www.alzforum.org/mutations/mapt), 35 of which are outside exon 10 and therefore present in both 3R and 4R *MAPT* transcripts. This is important as little 4R tau is expressed in iPSC-derived models without a splice-site mutation (Iovino et al., 2015; Sato et al., 2018; Sposito et al., 2015; Verheyen et al., 2018). Despite this limitation, iPSC-based models derived from patients with *MAPT* mutations have shown promise for studying molecular mechanisms underlying tauopathies, revealing altered transcriptomic signatures and tau accumulation and hyperphosphorylation (Gonzalez et al., 2018; Iovino et al., 2015; Jiang et al., 2018; Nakamura et al., 2019; Silva et al., 2016, 2020; Wray, 2017). The tau-V337M mutation is located in *MAPT* exon 12 and is expressed in iPSC-derived neural models. In the human brain it is associated with abundant neurofibrillary tangles in the frontal and temporal cortices and cingulate gyrus, and gliosis and atrophy throughout the hippocampus and amygdala (Spillantini et al., 1996; Spina et al., 2017; Sumi et al., 1992). In iPSC-derived induced neurons, tau-V337M can disrupt the axon initial segment and interfere with the regulation of neuronal activity and excitability (Sohn et al., 2019).

Brain organoids derived from iPSCs provide a new tool for disease modeling. These 3D tissues develop in culture along a trajectory that mimics key features of fetal brain development, and can be patterned towards specific brain regions, including those affected in tauopathy (Camp et al., 2015; Lancaster et al., 2017; Paşca et al., 2015; Velasco et al., 2019). Recent methodological advances have increased the replicability of organoid characteristics (Velasco et al., 2019; Yoon et al., 2019), rendering them suitable for large scale studies. Finally, increasing evidence that early alterations in brain development contribute to later manifestation of neurodegenerative diseases (Faravelli et al., 2020) have further encouraged the implementation of iPSC-based organoid systems to study FTD-tau.

Here, we report the characterization of over 6,000 cerebral organoids generated in a single centralized facility, derived from three tau-V337M mutation carriers and respective isogenic CRISPR corrected lines (Karch et al., 2019) (Table S1, Figure 1A). We show that tau-V337M organoids, relative to their isogenic controls, exhibit earlier expression of glutamatergic signaling pathways and synaptic genes, progressive accumulation of total tau and phosphorylated tau (P-tau S396), disruption of autophagy, and widespread changes in splicing, with later selective death of glutamatergic neurons. Finally, we demonstrate that glutamatergic cell death in the organoids can be pharmacologically blocked by apilimod treatment, which is predicted to target autophagy-lysosomal function and glutamate receptor recycling through PIKFYVE kinase inhibition (Cai et al., 2016; Shi et al., 2018).

**Figure 1.**
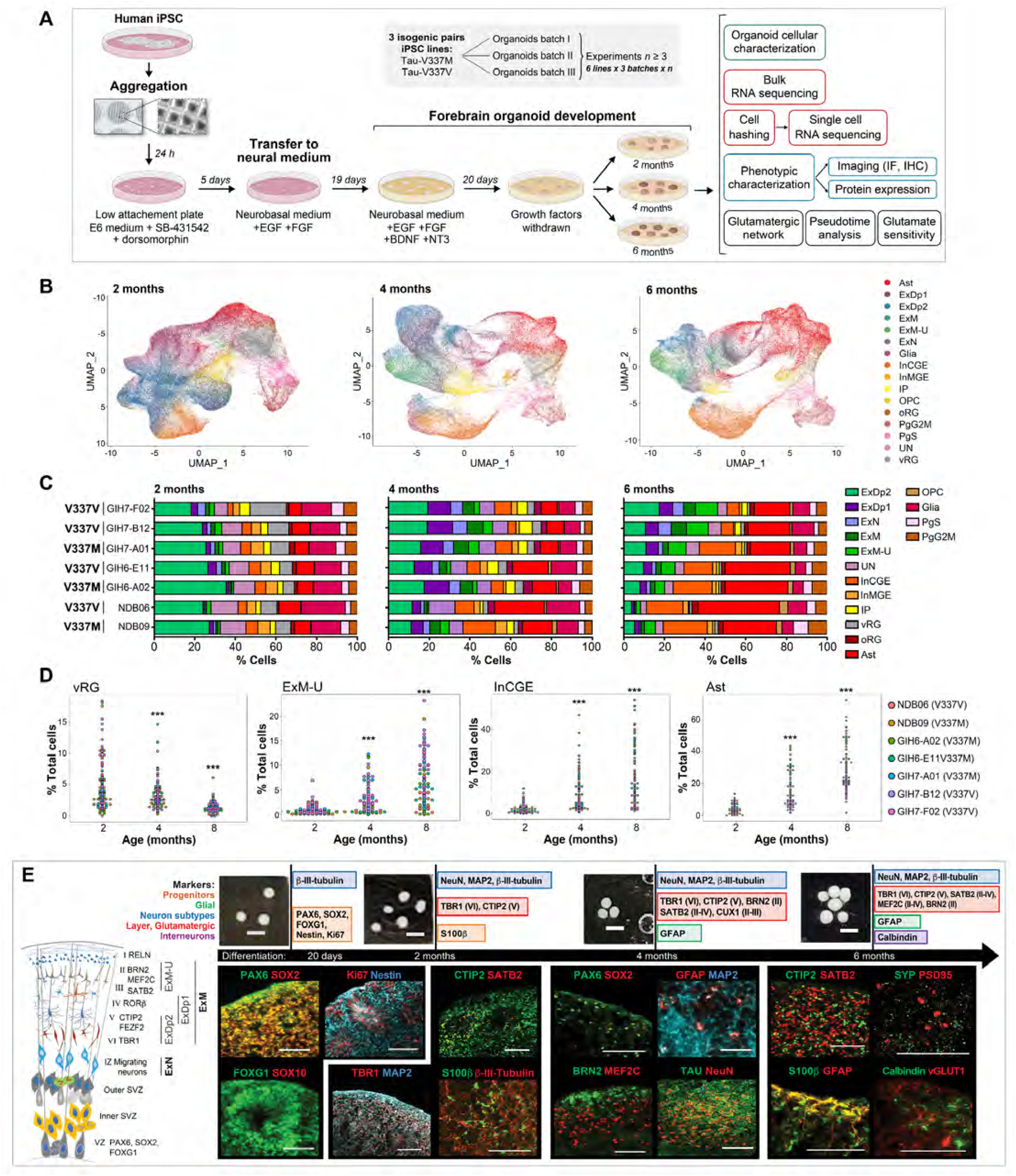
Forebrain Organoids Exhibit Similar Differentiation Patterns to Developing Fetal Brain. **(A)** Differentiation schematic and experimental summary. **(B)** UMAP reduction of scRNA-seq data at 2, 4 and 6 months, colored by cell type. Key: Ast-astrocytes, ExDp1-excitatory deep layer 1, ExDp2-excitatory deep layer 2, ExM-maturing excitatory, ExM-U-maturing excitatory upper enriched, ExN-migrating excitatory, Glia-unspecified glia/non-neuronal cells, InCGE-interneuron caudal ganglionic eminences, InMGE-interneuron medial ganglionic eminences, IP-intermediate progenitors, OPC-oligodendrocytes precursor cells, oRG-outer radial glia, PgG2M-cycling progenitors (G2/M phase), PgS-cycling progenitors (S phase), UN-unspecified neurons, vRG-ventricular radial glia. **(C)** Stacked bar plots of cell type proportions (%) per cell line at 2, 4, and 6 months. **(D)** Cell type proportions for individual organoids over time. *p<0.05, **p<0.01, ***p<0.001. **(E)** Schematic of organoid maturation and neural cell layering (left) and markers identification by IHC (right). Confirmation of dorsal forebrain progenitors at 20 days, by PAX6, SOX2, FOXG1, Nestin, proliferation marker Ki67 and absence of SOX10. 2 months: neurons express MAP2AB, β-III-tubulin and NeuN; deep layer glutamatergic neurons are observed by TBR1 and CTIP2/BCL11B staining, with few SATB2+ (upper layer) cells; S100β+ glia cells are detected. 4-6 months: few progenitor cells remain (SOX2, PAX6); Glutamatergic neurons of deep and upper layers are present, including BRN2, MEF2C expressing neurons and increased SATB2 cells; GFAP+ astrocytes and calbindin^+^ interneurons are present; Neurons robustly express tau; Expression of vGLUT1+ and pre- and post-synaptic markers, synaptophysin and PSD95 respectively. Scale bars 100 μm.

## RESULTS

### Single cell sequencing reveals selective loss of glutamatergic neurons in tau-V337M organoids

FTD-tau is associated with neuronal loss in the frontal and temporal cortices, particularly of glutamatergic neurons (Benussi et al., 2019). To determine whether glutamatergic neuronal loss is recapitulated in cerebral tau-V337M organoids we assessed changes in the proportions of different neural cell types over 6 months, a time period encompassing the generation of most cortical neuronal subtypes (Yoon et al., 2019).

Tau-V337M iPSC lines derived from three donors and corresponding isogenic corrected controls, totaling seven lines (Table S1, Karch et al., 2019), were differentiated into cerebral organoids (Yoon et al., 2019) and assessed at 2, 4 and 6 months using multiple phenotypic assays with three separate replicates (Figure 1A). We performed bulk RNA sequencing (RNAseq) on 239 individual organoids and single cell RNA sequencing (scRNA-seq) on 339 individual organoids using the 10x platform (sequencing over 370,000 cells across the samples), to pair global gene expression changes at high read depth with targeted transcriptomic profiling of individual cell types. By utilizing cell hashing, which uses antibody labeling to identify cell source (Stoeckius et al., 2018), we could assign cells to individual organoids, thus accounting for organoid-to-organoid variability and identify consistent effects across technical and biological replicates.

Following scRNA-seq data normalization, reduction and clustering, we conducted automated annotation with SingleR (Aran et al., 2019) using scRNA-seq expression data from human neocortex at gestational week (GW) 17-GW18 as a reference (Polioudakis et al., 2019), and identified 16 different cell types (Figure 1B-C). Organoid development followed the expected temporal changes (Lancaster et al., 2013; Velasco et al., 2019; Watanabe et al., 2017; Yoon et al., 2019), with decreased abundance of progenitor cells, including ventricular radial glial cells (vRGs), concomitant increase in neurogenesis, and later-onset gliogenesis (Figure 1B-D, Figure S1A). At 2 months, all organoids had similar cell composition, with a large proportion of deep layer excitatory (ExDp2) neurons (Figure 1B-C). Over time, the proportion of upper cortical layer enriched (ExM-U) neurons significantly increased, as did the proportion of inhibitory neurons, astrocytes and oligodendrocytes (all *p* < 2.2×10^-16^, Figure 1B-D). While all lines showed the same overall maturation pattern, the variability between lines and individual organoids increased over time (Figure 1C-D, Figure S1A), indicating donor line and clonal effects on long-term growth. There was no batch-specific effect on differentiation over three replicates assessed by UMAP (Figure S1B) and no apparent clustering effect due to the mutation at 2 or 4 months of differentiation (Figure S1C). Immunohistochemical (IHC) analysis of organoid sections confirmed the progressive maturation of organoids observed by scRNA-seq (Figure 1E, Figure S1D-E). Consistent with the inside-out order of cortical neurogenesis, markers for deep cortical layers VI (TBR1) and V (CTIP2/BCL11B) were present from 2 months of differentiation, and markers for upper layers IV-II (SATB2, MEF2C, BRN2) were robustly present from 4-6 months, along with synaptic markers (Figure 1E).

We examined whether tau-V337M impacted the survival of excitatory neurons, as observed in FTD brain (Benussi et al., 2019). While there were no differences in neuronal proportions at 2 or 4 months in any excitatory neuron category, there was a significant reduction in deep layer excitatory neurons (ExDp2, *p* = 0.004), maturing excitatory neurons (ExM, *p* = 0.003) and the proportion of newborn neurons (ExN, *p* = 0.0001) in 6-month tau-V337M organoids relative to V337V (Figure 2A). These differences were validated by IHC analysis that showed reductions in NeuN+ neurons at 4 and 6 months (Figure 2B-C) and MAP2+ neurons at 6 months (Figure S2A-B). In contrast, there was no such effect on interneurons or astrocytes (Figure S2C). These results demonstrate that tau-V337M cerebral organoids recapitulate aspects of selective vulnerability of excitatory neurons observed in the human brain (Benussi et al., 2019).

**Figure 2.**
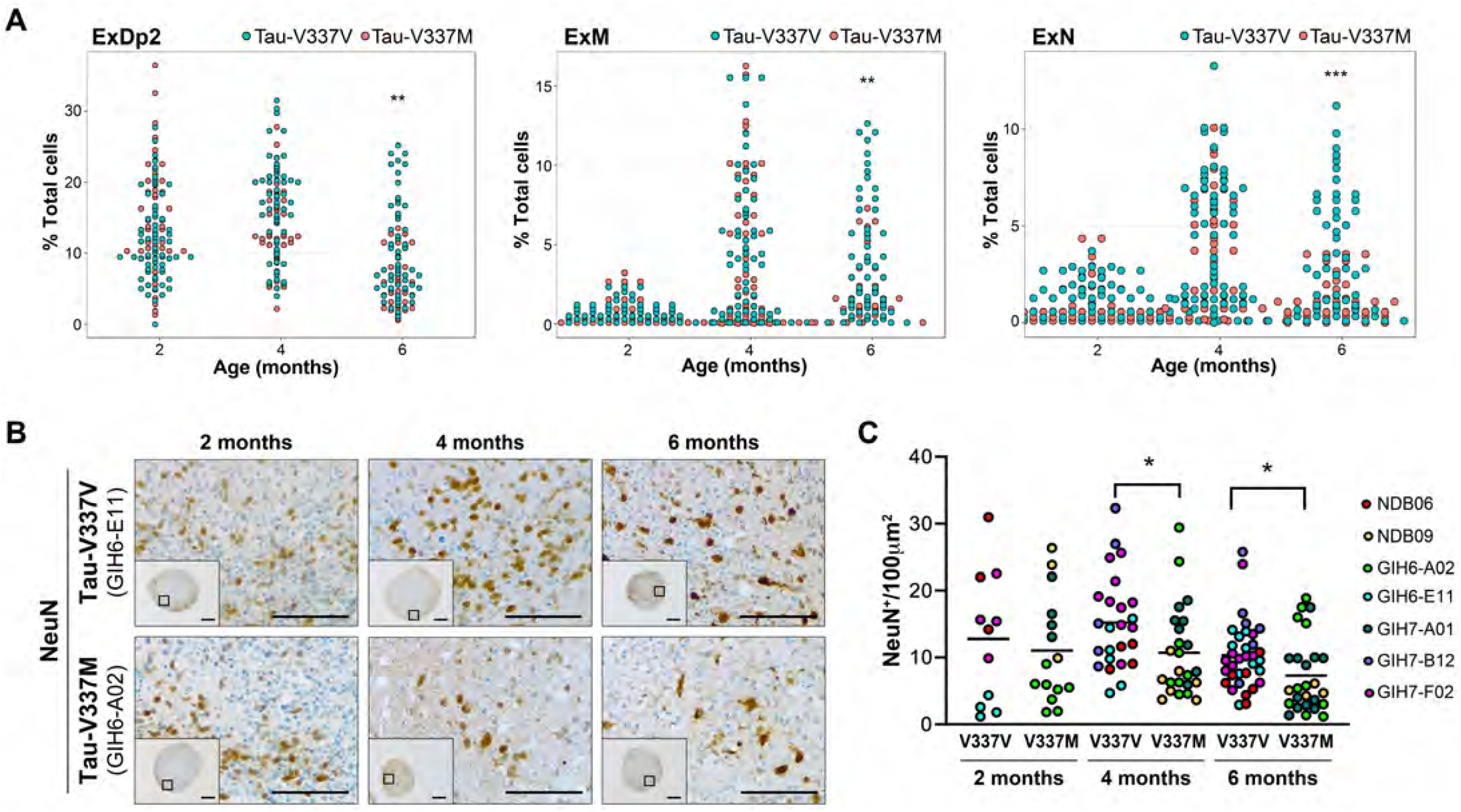
Tau-V337M Organoids Exhibit Neuronal Loss Phenotypes. **(A)** Proportion of deep layer glutamatergic neurons (ExDp2, ExM and ExN) per organoid at 2, 4 and 6 months. \*\**p*<0.01, \*\*\**p*<0.001. **(B-C)** Imaging and quantification of neuronal density by NeuN+ at 2, 4 and 6 months, in tau-V337M and isogenic V337V organoids. *p<0.05. Scale bars are 250 μm (inserts) and 50 μm.

### V337M organoids exhibit increased tau and P-tau and degeneration of deep cortical layers

In tau-V337M human brain, neurofibrillary tangles follow a ‘tram-track’ pattern with stronger staining in cortical layers II and V (Spillantini et al., 1996; Spina et al., 2017) that appear particularly vulnerable to tau accumulation and selective neuronal loss. Since tau accumulation and disruption of autophagy-lysosomal function are key FTD hallmarks (Bain et al., 2019; Piras et al., 2016; Wang et al., 2009), we examined these characteristics in mutant versus isogenic control organoids over time.

Tau-V337M organoids showed a significant increase in total tau and the P-tau S396/total tau ratio between 4 and 6 months compared to isogenic tau-V337V organoids (Figure 3A-B). No genotype-dependent differences were observed at 2 months of differentiation. The increased burden of tau with age mirrors prior observations in 2D iPSC-derived neuronal models (Ehrlich et al., 2015; Fong et al., 2013; Iovino et al., 2015; Silva et al., 2016, 2020; Wren et al., 2015) and in human brain from V337M carriers (Spillantini et al., 1996; Spina et al., 2017).

**Figure 3.**
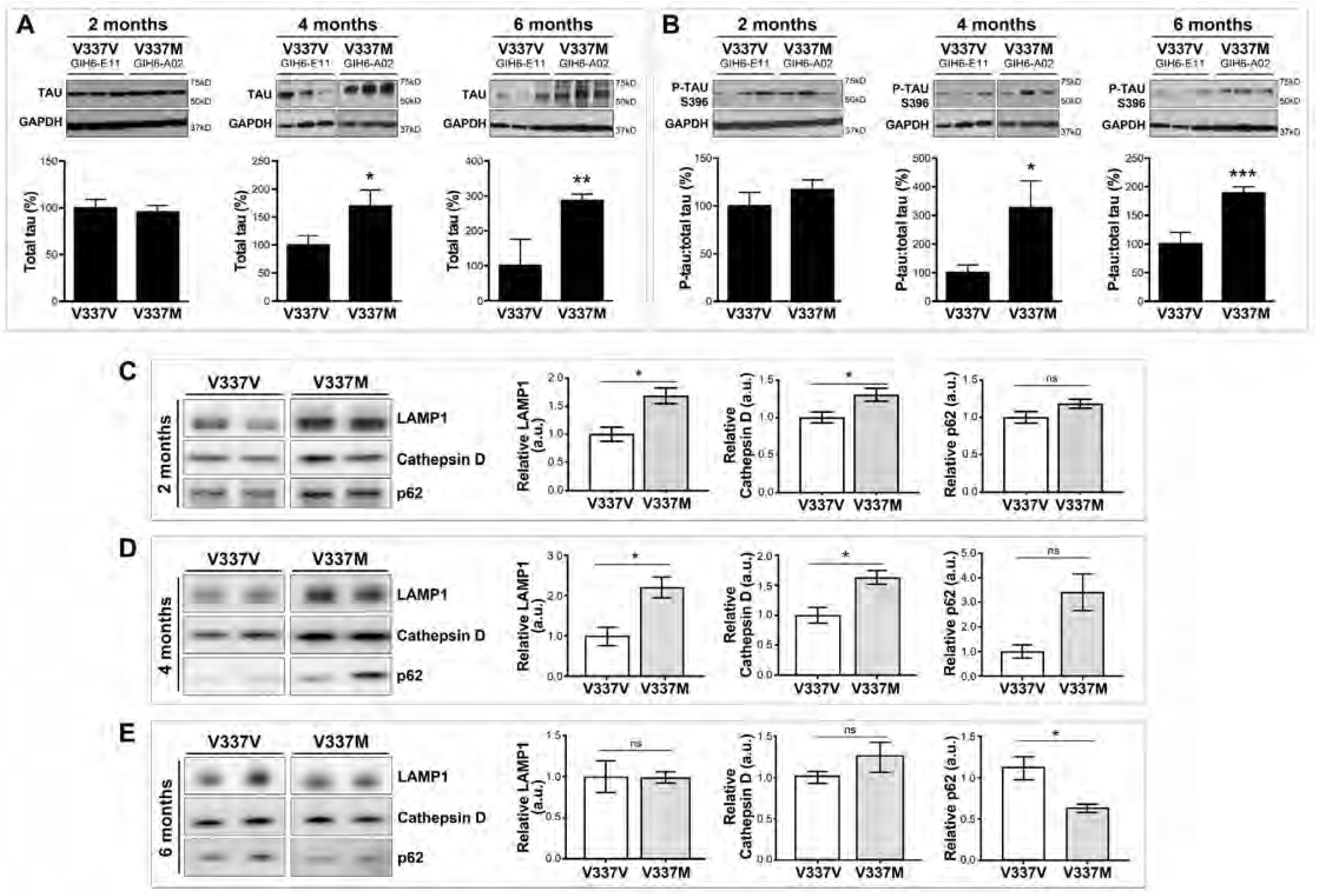
Tau-V337M Organoids Reveal Progressive Tau Accumulation and Early Autophagy Disruption. **(A-B)** Western blot and densitometry quantification of total tau **(A)** and P-tau S396 **(B)** levels in tau-V337M and isogenic V337V organoids at 2, 4 and 6 months. Bars represent tau densitometry in mutant organoids (%) relative to isogenic controls, ±SEM. \**p* ≤ 0.01, \*\**p* ≤ 0.001. *n* = 9 per group from 3 independent experiments. **(C-E)** Western blot and densitometry quantification of autophagy-lysosomal pathway markers in tau-V337M and isogenic organoids at **(C)** 2 months, **(D)** 4 months and **(E)** 6 months of differentiation. Graph bars represent ± SEM. \**p* ≤ 0.01, ns/ non-significant. *n* = 3 organoids per group.

To examine whether the tau-V337M mutation was associated with disruption of autophagy, we measured LAMP 1 (lysosomal associated membrane protein 1), Cathepsin D and p62 levels in organoid lysates (Figure 3C-E). LAMP1 and Cathepsin D were significantly increased in the tau-V337M organoids compared to the isogenic controls at 2 months (Figure 3C), and p62 was significantly reduced at 6 months (Figure 3D). These findings show that iPSC-organoids phenocopy autophagy-lysosomal and proteostatic dysfunction seen in FTD brain, and that these may be triggered by the *MAPT* mutation early in disease development.

Analysis of the bulk transcriptomic data indicated that total *MAPT* mRNA expression was higher in tau-V337M organoids compared to V337V at 2 months, however this effect was no longer observed at 4 months and was reversed at 6 months (Figure S3A), potentially reflecting neuron loss at this time point (Figure 2A-C). Hence, increased tau protein at 6 months (Figure 3A-B) is likely due to tau accumulation and not increased expression. The scRNA-seq data revealed that *MAPT* expression was highest in excitatory glutamatergic neurons (Figure 4A-C, Figure S3B), particularly in neurons of cortical layers VI-V (ExDp2; Figure 4C). Moreover, *MAPT* expression was significantly higher in tau-V337M ExDp2 neurons compared to V337V controls at every time point (2 months *p* = 6.49×10^-23^, 4 months *p* = 1.88×10^-18^, 6 months *p* = 3.27×10^-08^), and in V337M ExDp1 and ExM-U at 4 months (ExDp1 *p* = 1.16×10^-07^, ExM-U *p* = 2.82×10^-04^; Figure 4C). Notably, this links high expression of *MAPT*, and of mutated *MAPT*, to the later selective vulnerability of the same neuronal subtypes.

**Figure 4.**
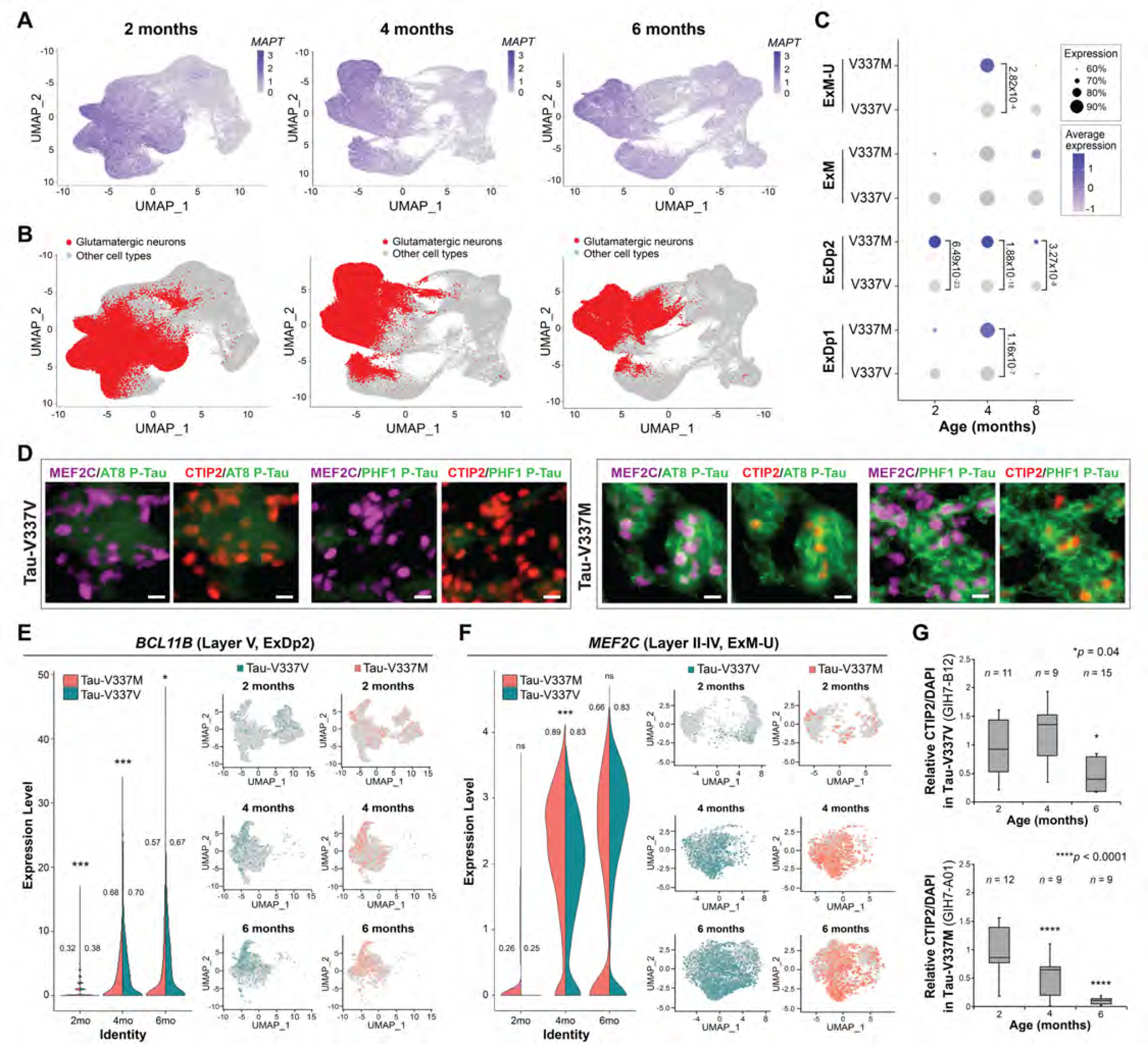
Tau-V337M Organoids Reveal Loss of Deep-Layer Glutamatergic Neurons. **(A-B)** Expression of *MAPT* **(A)** and glutamatergic neuronal subtypes (ExDp1, ExDp2, ExM and ExM-U) **(B)** projected onto scRNA-seq UMAP plots at 2, 4 and 6 months. **(C)** Proportion of *MAPT*-expressing glutamatergic neuronal subtypes over time, by mutation. Expression scaled within each time point. Dot size = proportion of cells expressing *MAPT*, depth of color = *MAPT* expression level. Values = differential gene expression p-value adjusted by MAST general linear model comparisons of differential expression. **(D)** IHC of tau-V337M (right) and isogenic control (left) 6-month organoids illustrates CTIP2/BCL11B+ and MEF2C+ neurons with P-tau S202/T205 (AT8) and P-tau S396/S404 (PHF1) staining. Scale bar 10 μm. **(E-F)** Proportion of V337M and V337V ExDp2 neurons expressing layer V marker CTIP2/*BCL11B* **(E)** or ExM-U neurons expressing layer II-IV marker *MEF2C* **(F)** at each time point; values = proportion of cells expressing each gene, \**p* < 0.05, \*\**p* < 0.01, \*\*\**p* < 0.001 between V337M and V337V neurons. UMAPs of gene expression in ExDp2 neurons **(E)** or ExM-U neurons **(F)** at each time point. **(G)** Time-course IHC quantitative analysis of CTIP2+ neurons at 2, 4 and 6 months. Box plots illustrate distribution of CTIP2/BCL11B+ neurons normalized to DAPI from ≥ 3 different organoids from 3 separately generated organoid batches. *n* = number of organoids, *p < 0.05, **** p < 0.0001.

Given that cortical layer II and V neurons are vulnerable populations in FTD (Ghetti et al., 2015), we investigated whether this selective vulnerability was observed in the organoid model. IHC analysis revealed P-Tau S202/T205 (AT8) and P-Tau S396/S404 (PHF1), modifications seen with tau pathology in V337M patients (Spillantini et al., 1996), in cells expressing MEF2C (which with maturation is expressed predominantly in layers II-IV (Trevino et al., 2020)) and the deep layer V marker *BCL11B/CTIP2* (Figure 4D), corroborating the tram-track tau pathology observed in V337M human brain tissue (Spillantini et al., 1996; Spina et al., 2017). Analysis of the scRNA-seq data showed significantly fewer glutamatergic neurons expressing *MEF2C* and *BCL11B* in V337M vs. V337V organoids at 6 months, despite similar expression at 2 and 4 months (Figure 4E-F). A similar pattern of loss was observed by IHC: V337M organoids exhibited a significant decrease in CTIP2+ neurons at 4 and 6 months relative to 2 months (Figure 4G). Hence, mutant tau-V337M organoids replicate the cortical layer vulnerability observed in human FTD post-mortem brain; increased expression of *MAPT* and accumulation of P-tau/tau in these neuronal subpopulations are likely contributors to this vulnerability.

### Early neuronal maturation and upregulation of synaptic signaling pathways in mutant glutamatergic neurons

To determine early changes underlying neuronal dysfunction prior to tau aggregation and neurodegeneration, we conducted Gene Set Enrichment Analysis (GSEA) on differentially expressed genes at 2, 4, and 6 months in the bulk RNA-seq data (Figure S4A-F). In mutant organoids, we observed upregulation of numerous synapse-related pathways compared to isogenic controls at 2 months (Figure S4A, bold text), but downregulation between 2 and 6 months (Figure S4B). Further inspection of specific genes within these pathways emphasized glutamatergic genes and revealed that both ionotropic and metabotropic glutamatergic receptors exhibited this non-monotonic pattern of early upregulation and late downregulation in mutant compared to control, including *GRM5* (*p* = 1.25×10^-04^, log2 fold change (FC) = 1.08), *GRIN1* (*p* = 3.89×10^-03^, FC = 0.47), and *GRM4* (*p* = 6.9×10^-03^, FC = 0.64) (Figure 5A, Figure S4G).

**Figure 5.**
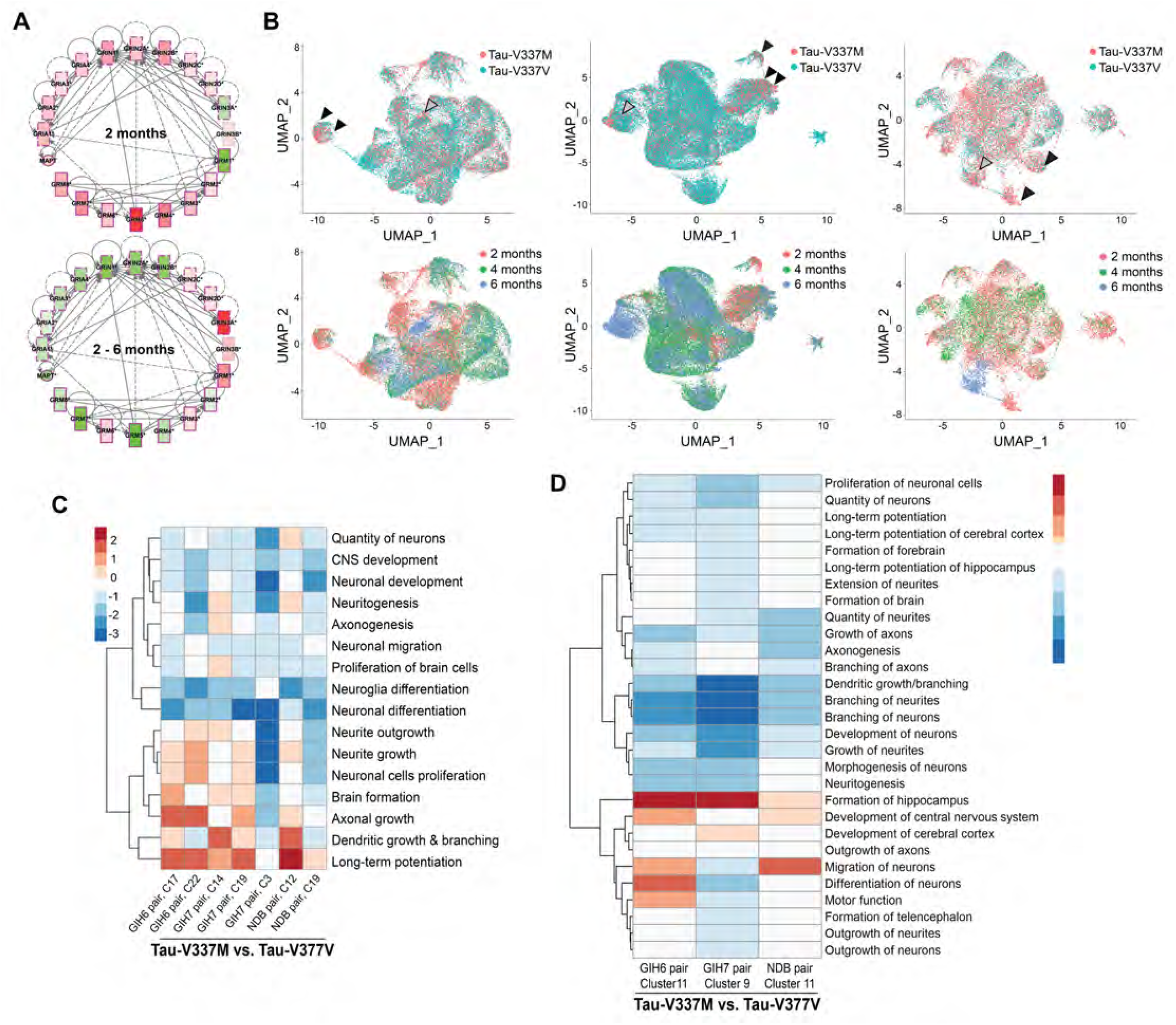
Tau-V337M Organoids Exhibit Early Neuronal Maturation and Upregulation of Synaptic Signaling Pathways. **(A)** Expression and connectivity of glutamatergic receptor genes and *MAPT* at 2 months and 2-6 months. Red = upregulation in tau-V337M organoids compared to isogenic controls, green = downregulation. Depth of color = extent of expression fold-change. **(B)** UMAP reduction of glutamatergic neurons for each organoid isogenic pair, colored by mutation (upper) and age (lower). 2-month V337M-enriched clusters indicated by black arrowheads, and 6-month V337M-enriched clusters indicated by grey arrowheads. **(C-D)** Z-scores for enriched pathways derived from Ingenuity pathway analysis for V337M-enriched 2-month **(C)** and 6-month **(D)** glutamatergic neuronal clusters (C#) for three isogenic pairs.

To further explore altered synaptic signaling pathways and specific loss of glutamatergic neurons over time, we examined excitatory neuronal populations in the scRNA-seq data (ExDp2, ExDp1, ExM, ExM-U). For each isogenic pair of lines, excitatory neurons were re-clustered and V337M-enriched clusters were identified (FC > 1.5; Figure 5B) and assessed for differential gene expression. Tau-V337M enriched clusters of 2-month-old excitatory neurons showed downregulation of pathways such as *Differentiation of Neuroglia* (z-score = ^-^0.274 - ^-^2.175), *p* = 3.78×10^-08^- 4.67×10^-11^) and *Differentiation of Neurons* (z-score = ^-^1.131 - ^-^2.606, *p* = 1.74×10^-09^-1.39×10^-20^) in all three donor lines (Figure 5C). Consistent with the bulk sequencing data, pathways associated with neuronal maturation and synaptic signaling, such as *Growth of Axons* (z-score = 0.239-1.482, *p* = 9×10^-06^-2.3×10^-13^), *Dendritic Growth/Branching* (z-score = 0.465-1.608, *p* = 2.37×10^-09^-5.05×10^-07^), and *Long-term Potentiation* (z-score = 0.102-2.775, *p* = 4.1×10^-11^-6.91×10^-05^; Figure 5C) were upregulated at 2 months. These pathways were downregulated at 6 months in V337M-enriched clusters (Figure 5D).

We next carried out pseudotime analysis using Monocle3 (Qiu et al., 2017; Trapnell et al., 2014) to order cells by their differentiation state. We validated its ability to separate and order cell types as anticipated, with deep cortical layer neurons (ExDp2) tending to occur earlier than upper layer neurons (ExM, ExM-U) (Figure S5A-E). V337M-enriched modules occurred principally in early pseudotime encompassing primarily ExDp2 neurons, indicating early aberrant gene expression networks (Figure S5F). Of the top 3,000 most variable genes across all excitatory neurons, 391 had significantly different trajectories in mutant neurons compared to isogenic controls; gene ontology (GO) analysis revealed significant enrichment of glutamatergic signaling pathways, including *Glutamate Binding* (*p* = 0.02), *Glutamatergic Synapse* (*p* = 0.008), *Clathrin-Sculpted Glutamate Transport Vesicle* (*p* = 0.009) and *Glutamate Neurotransmitter Release Cycle* (*p* = 0.05). We then clustered the significantly different genes across pseudotime to identify those with similar expression trajectories (Figure 6A). Of the 27 genes associated with enriched glutamatergic signaling GO pathways, 11 were present in cluster 2, including the neuronal splicing regulator ELAVL4; *GRIA2* and *MAPT* were also present in this cluster. (Figure 6A-C). This cluster was characterized by an earlier upregulation and expression of genes across pseudotime in tau-V337M neurons relative to V337V that typically converged at later pseudotime points (Figure 6B-C). GO enrichment of genes within cluster 2 showed the most significant enrichment for *Neuron Projection* (*p* = 3.63×10^-15^), *Synapse* (*p* = 1.34×10^-13^) and *Axon* (*p* = 2.14×10^-13^) cellular compartments, and *Nervous System Development* (*p* = 5.35×10^-13^) and *Synapse Organization* (*p* = 7.1×10^-12^) molecular functions.

**Figure 6.**
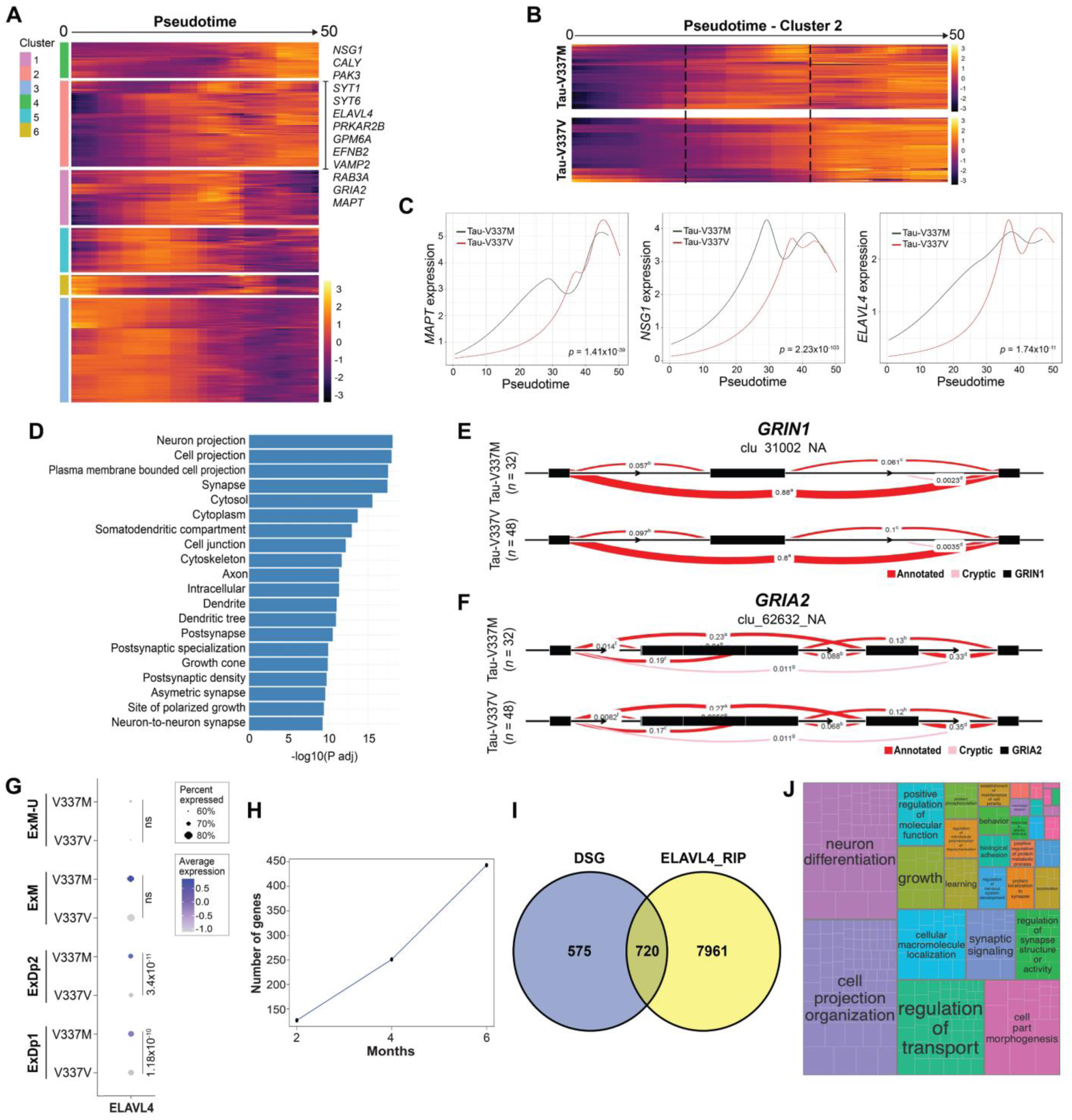
Accelerated Glutamatergic Gene Expression Precedes Aberrant Splicing in V337M Neurons. **(A)** Expression of genes with significantly differential trajectories over pseudotime in tau-V337M glutamatergic neurons. Patterns of expression over pseudotime are grouped by unsupervised hierarchical clustering and centered across genes. Low expression = purple, high expression = yellow. Genes present in significantly enriched glutamatergic signaling pathways are highlighted in cluster 2. **(B)** Comparison of cluster 2 pseudotime trajectories in V337M and V337V glutamatergic neurons. Dashed lines highlight the central region of pseudotime where gene expression differs between mutant and control cells. **(C)** Trajectories of average *MAPT*, *NSG1* and *ELAVL4* expression in V337M and V337V glutamatergic neurons over pseudotime. Adjusted *p*-value for statistical comparison of trajectories is shown in the bottom right corner of each plot. **(D)** GO pathways enriched for differentially spliced genes between 6-month V337M and V337V organoids. Bars indicate adjusted-log10 *p*-value. **(E-F)** Differentially spliced intron clusters within glutamatergic receptor genes *GRIN1* **(E)** and *GRIA2* **(F).** Exons = black boxes. Red band thickness and inserted values represent proportion of spliced exon-exon pairs. **(G)** Expression of *ELAVL4* in V337M and V337V glutamatergic neurons. Dot size = proportion of cells expressing *MAPT*, depth of color = *MAPT* expression level. Values = differential gene expression *p*-value adjusted by MAST general linear model comparisons of differential expression. **(H)** Number of ELAVL4-bound differentially spliced genes over time. **(I)** Proportion of overlap between number of differentially spliced genes in tau-V337M organoids and genes bound to ELAVL4 by RIP analysis. **(J)** Semantic analysis of significant GO pathways enriched for ELAVL4-bound differentially spliced genes.

### Splicing changes in synaptic signaling pathways accompany degenerative changes in mutant organoids

Splicing analysis using Leafcutter (Li et al., 2018; Figure 5D-F, Figure S5G) showed a dramatic increase in the number of differentially spliced junctions in genes of tau-V337M compared to tau-V337V organoids, from 190 at 2 months to over 5,000 at 6 months (Figure S5G). GO analysis of the differentially spliced genes at 6 months revealed a significant enrichment in several synaptic pathways as well as cellular projections (Figure 6D). Among these were exon 5 (chr9:137145902-13749009) of the NMDA receptor gene *GRIN1* (log likelihood ratio (loglr) = 12.6, *p* = 2.76×10^-3^), which was more frequently excluded in V337M organoids compared to V337V (differential percent spliced in (dPSI) = −0.06); Figure 6E), and exon 14 (chr4:157360143-157361538) of the AMPA receptor gene *GRIA2* (loglr = 27.18,*p* = 1.96×10^-6^), which was more frequently included in V337M organoids (dPSI = 0.038; Figure 6F). Therefore, while the V337M mutation was not found to impact *MAPT* splicing, it did impact the regulation of splicing in pathways associated with neuronal and synaptic maturation that may contribute to the glutamatergic neuronal vulnerability observed at 6 months.

*ELAVL4*, which showed altered expression over pseudotime, and was present in cluster 2 along with aberrantly expressed *MAPT* and glutamatergic genes (Figure 6B-C), is a neuronal specific RNA binding protein (RBP) associated with neural development, synaptic plasticity, splicing, glutamate receptor activation and glutamate levels, as well as multiple neurological diseases (Bolognani et al., 2009; Ince-dunn et al., 2012; Tiruchinapalli et al., 2008; Bronicki and Jasmin, 2013; Zhou et al., 2011; Zhu et al., 2008)). Expression of *ELAVL4* was enriched in lower cortical layers of tau-V337M glutamatergic neurons (ExDp2, ExDp1; Figure 6G), consistent with different cortical layer expression of *ELAVL* family genes observed in human brain (Ravanidis et al., 2018). Given its previously defined roles, *ELAVL4* may be relevant to the susceptibility of this neuronal population.

To further assess the potential contribution of *ELAVL4* expression to altered splicing regulation of synaptic and glutamatergic genes in tau-V337M organoids, we determined whether there was a significant intersection between our observed differentially spliced genes and RNA immunoprecipitation sequencing (RIP-seq) data for ELAVL binding in the human brain (Scheckel et al., 2016; Tebaldi et al., 2018). The proportion of transcripts that overlapped with ELAVL4 cargo increased greater than 3-fold with age, resulting in over 50% of differentially spliced genes as potential ELAVL4 cargo (Figure 6H-I). GO enrichment of ELAVL4-bound genes revealed processes associated with *RNA splicing* (*p* = 2.4×10^-08^, FDR 0.2.2X10^-06^) and *Neuron Differentiation* (*p* = 2.1×10^-24^, FDR = 1.4×10^-20^), and semantic analysis of all significant GO categories (FDR <0.1) indicated enrichment of pathways associated with neuron differentiation, cell projection organization and other categories associated with neuronal function (Figure 6J), suggesting that aberrant *ELAVL4* expression is likely to impact these neuronal functions in tau-V337M organoids.

### Excitotoxic stress vulnerability and selective neuronal death in V337M organoids is reversed by apilimod

Since our transcriptomic analyses indicated that tau-V337M organoids display early increased expression of glutamatergic signaling components at 2 months followed by selective glutamatergic neuronal death at 6 months, we examined whether neurons in tau-V337M organoids were more vulnerable to excess glutamate stimulation than isogenic tau-V337V controls.

To avoid confounding factors such as the formation of new neurons (Shi et al., 2019) and to better model advanced disease, we used both 2 and 12 month organoids and established a method for longitudinal tracking of individual neurons by transduction with a *Synapsin::GFP* lentivirus and immobilization of each organoid in matrigel (Figure 7A). Following initial imaging to establish a baseline neuron number per organoid, 5 mM glutamate was added to the culture media at one, two, four, and six days. Organoids were imaged daily during this period to track individual neurons over time. In the absence of excess glutamate in the media, neurons in control and tau-V337M organoids survived equally well (Figure 7B). In contrast, repeated treatment with 5 mM glutamate triggered a more rapid and significant loss of tau-V337M neurons relative to that observed in isogenic controls (*p* < 0.0001, Figure 7C-D). Thus, neurons in tau-V337M organoids were more sensitive to glutamate-induced toxicity than neurons in control organoids.

**Figure 7.**
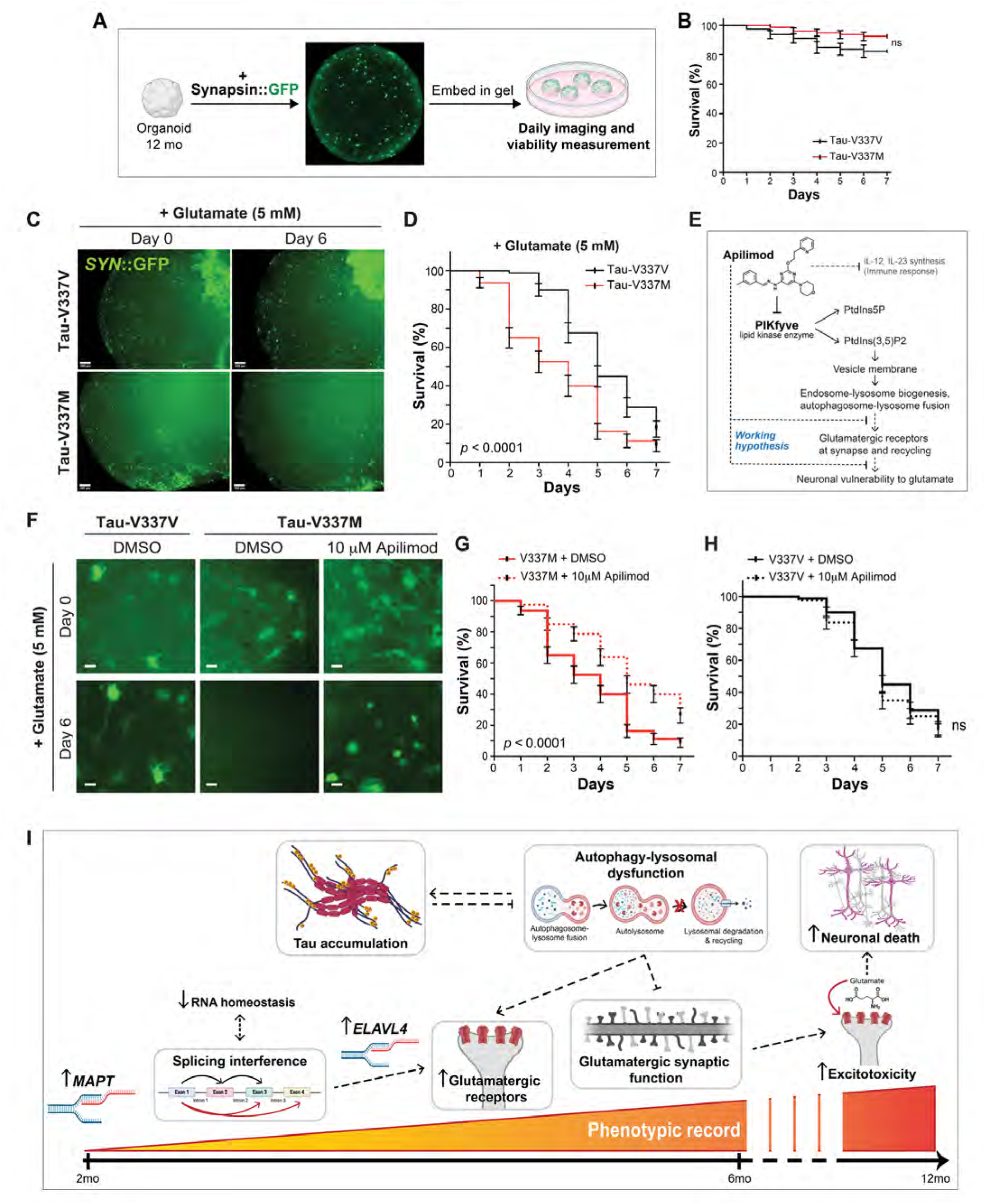
Susceptibility to Glutamate Excitotoxicity in Tau-V337M Organoids Is Reversed by Apilimod. **(A)** Method for tracking neuronal survival in cerebral organoids using longitudinal imaging. **(B)** Survival of *Syn*::GFP+ neurons in 12-month tau-V337M and isogenic V337V organoids without glutamate treatment. *n* = 80 neurons tracked from 5 individual organoids per group. ns/not significant. **(C)** Images of tau-V337M and isogenic V337V organoids treated with 5 mM glutamate. Neurons were labeled with a lentivirus encoding *Syn*::GFP. Scale bars 100 μm. **(D)** Survival of *Syn*::GFP+ neurons in tau-V337M and isogenic V337V 12-month organoids with glutamate treatment. *n* = 80 neurons tracked from 5 individual organoids per group. \*\*\*\**p* < 0.0001. **(E)** Schematic of proposed mode of action for apilimod on neuronal vulnerability. **(F)** Images of 12-month tau-V337M and isogenic V337V organoids treated with 5 mM glutamate and DMSO or 10 μM apilimod. Neurons labeled with *Syn*::GFP. Scale bars 10 μm. **(G-H)** Survival of *Syn*::GFP+ neurons in tau-V337M **(G)** organoids and isogenic V337V **(H)** organoids with glutamate treatment and DMSO or apilimod. *n* = 80 neurons tracked from 5 individual organoids per group. \*\*\*\**p* < 0.0001, ns/not significant. **(I)** Working model for temporal molecular changes contributing to glutamatergic neuronal vulnerability and death.

PIKFYVE is a lipid kinase that regulates endolysosomal trafficking by converting phosphatidylinositol-3-phosphate (PI3P) into phosphatidylinositol-3,5-bisphosphate (PI3,5P2; Marat and Haucke, 2016) (Figure 7E). Previous studies have shown that after NMDA-induced endocytosis of glutamate receptors, PIKFYVE activity promotes RAB-dependent recycling of the receptors back to the cell surface (Seebohm et al., 2012). Small molecule inhibition of PIKFYVE (e.g., apilimod) can prevent recycling of receptors back to postsynaptic densities and protect against NMDA-induced excitotoxicity *in vitro* and *in vivo* (Shi et al., 2018; Figure 7E). Based on this evidence, we hypothesized that PIKFYVE inhibition might rescue the glutamate-induced neuronal loss observed in tau-V337M organoids (Figure 7E). To test this, we performed longitudinal tracking of *Synapsin*::GFP-labeled neurons with or without 10 μM apilimod treatment. Apilimod fully rescued the survival deficit specific to neurons in tau-V337M cortical organoids in response to excess glutamate (p<0.0001), yet had no effect on the survival of control tau-V337V neurons under the same conditions (Figure 7F-H). Altogether, our results reveal that early deficits in the glutamatergic neuronal network can be detected in FTD patient iPSC-derived cortical organoids, and that vulnerability to excitotoxicity can be pharmacologically blocked.

## DISCUSSION

We have characterized a 3D cerebral organoid model of FTD, using iPSC lines from three donors carrying the tau-V337M mutation and isogenic corrected controls, in order to identify the temporal changes leading to neurodegeneration. This model exhibits progressive accumulation of total tau and P-tau, early impaired autophagy function and later, specific loss of glutamatergic neurons. Significant upregulation of glutamatergic signaling pathways and increased autophagy-lysosomal pathway markers, as well as splicing changes, are seen early in mutant neurons. We identified early upregulated *ELAVL4* expression that may contribute to impaired splicing and accelerated glutamatergic synaptic gene expression. Additionally, we show that these abnormalities lead to neuronal dysfunction and vulnerability to excitotoxicity. Hence, using this human organoid model system, we have uncovered a time course of molecular changes in critical pathways associated with a *MAPT* mutation that ultimately result in specific, preferentially deep layer, glutamatergic neuronal death.

Given that organoid models do not accurately reflect adult aging, it is interesting that we observed neurodegeneration. However, this may reflect increased stress on a vulnerable neuronal population, because the organoid environment can itself induce cell stress (Bhaduri et al., 2020), which may contribute to accelerated neuronal death. We show that expression of tau-V337M resulted in progressive accumulation of both total tau and P-tau S396 as organoids aged, consistent with human FTD brain observations (Shiarli et al., 2006; Spillantini et al., 1996; Sumi et al., 1992). Furthermore, autophagy-lysosomal markers were increased in tau-V337M organoids, indicative of impaired function (Wang et al., 2009), which corroborates reports of increased LAMP1 and accumulation of autophagic vesicles in corticobasal degeneration (CBD) and progressive supranuclear palsy (PSP) (Piras et al., 2016), and other models of tauopathy (Lin et al., 2003; Nixon et al., 2005; Yu et al., 2004). As impairment of the autophagy-lysosomal system has been identified as a key contributor to the pathological accumulation of tau (Bain et al., 2019; Bendiske and Bahr, 2003; Wang et al., 2009), it is likely that pathway dysfunction associated with mutant tau further contributes to the observed tau accumulation. Together, these data indicate that tau-V337M neurons exhibit early and progressive dysfunction in important cellular pathways preceding overt tau pathology such as neurofibrillary tangles. These findings link early molecular changes in cellular function to the manifestation of neurodegenerative disease in later life.

Interestingly, in the cortex of tau-V337M patients, tau neurofibrillary tangles occur primarily in a tram-track pattern, corresponding to cortical layers II and V (Spillantini et al., 1996; Spina et al., 2017), indicating a selective vulnerability of these neurons. However, the reason for this vulnerability is unknown. We found that *MAPT* expression was highest in deep layer glutamatergic excitatory neurons encompassing cortical layer V, consistent with the vulnerability of these neurons to tau accumulation seen in human brain, and subsequent neurodegeneration of layer V neuronal subsets such as von Economo neurons (Lin et al., 2019). Indeed, we observe loss of layer II and V neurons at 6 months in tau-V337M organoids, indicating that higher *MAPT* expression may underlie selective vulnerability of these cells.

Our studies point to major and early alterations in glutamatergic signaling pathways associated with the *MAPT* mutation. Glutamate is the main excitatory neurotransmitter, and glutamate excitotoxicity is the predominant cause of cerebral damage (Benussi et al., 2019), which can induce neurodegeneration (Bowie, 2008). Transcranial magnetic stimulation studies have revealed an impairment in glutamatergic circuits in both sporadic and inherited FTD (Borroni et al., 2018). Antibodies against the AMPA receptor GluA3 were identified in the serum and CSF of FTD patients (Borroni et al., 2017), and reduced availability of the metabotropic receptor mGluR5 was observed in behavioral variant FTD (Leuzy et al., 2016). Animal models also suggest impaired glutamatergic signaling: AMPA and NMDA receptor hypofunction was identified in the ventral striatum and insular cortex of tau-V337M mice (Warmus et al., 2014). Given that much of the human and mouse data suggest a reduction or an impairment in glutamatergic signaling, it initially seemed inconsistent that we identified an early upregulation of this pathway. However, we demonstrate that this was followed by increased susceptibility to excitotoxicity in tau-V337M organoids and a reduction in these pathways as cells become dysfunctional and die. Indeed, there is evidence of increased seizure susceptibility in FTD patients early in disease (Beagle et al., 2017). Electrophysiological recordings from tau-P301L, tau-N279K and tau-V337M iPSC-derived neurons in 2D culture have shown earlier synaptic maturation and increased excitability compared to controls (Iovino et al., 2015; Sohn et al., 2019). These early changes in synaptic maturation and glutamatergic signaling are therefore likely to predispose neurons to dysfunction and excitotoxicity, resulting in later synaptic loss and neurodegeneration, and may be common across different *MAPT* mutations.

There are several potential mechanisms by which tau-V337M may prematurely enhance synaptic maturation and glutamatergic signaling. One is by altered regulation of alternative splicing, which is crucial for synaptic maturation (Weyn-Vanhentenryck et al., 2018). We observe early splicing dysregulation in tau-V337M organoids in genes that converge on synaptic signaling pathways. It has also been hypothesized that mutant tau mislocalizes to the somatodendritic compartment, where it alters RBP function (Vanderweyde et al., 2016). In the tau-P301S mouse, aggregation of RBPs with tau, and resulting alterations in splicing, converge on pathways associated with synaptic transmission (Apicco et al., 2019), similar to the effect we observed in tau-V337M organoids. These mice exhibit increased inclusion of exon 14 of the AMPA receptor subunit *GRIA2* (Apicco et al., 2019), which we also observed in tau-V337M organoids. This splicing change is consistent with an excitatory phenotype, because expression of this exon leads to slower desensitization of AMPA receptors, increasing susceptibility to excitotoxicity (Koike et al., 2000; Pei et al., 2009). We also identified increased exclusion of *GRIN1* exon 5 in tau-V337M organoids, which has been associated with the overproduction of excitatory synapses in layer V pyramidal neurons in adult mice that in turn increases seizure susceptibility (Liu et al., 2019), and may contribute to the cortical layer vulnerability observed in our model and in FTD brain.

The neuronal-specific RNA-binding gene *ELAVL4* is selectively expressed in vulnerable neuronal populations in tau-V337M organoids, consistent with its observed expression in cortical glutamatergic neurons of layers V and VI in brain (Bronicki and Jasmin, 2013; DeBoer et al., 2014). Notably, *ELAVL4* expression decreases self-renewal of neural progenitor cells and promotes neuronal differentiation (Akamatsu et al., 2005; DeBoer et al., 2014), matching our observation of accelerated synaptic maturation in tau-V337M organoids. *ELAVL4* regulates RNA splicing, glutamate levels and neuronal excitability (Ince-dunn et al., 2012), and it binds to the metabotropic glutamatergic receptor *GRM5* and multiple genes within the LTP pathway (Bolognani et al., 2009), which were upregulated in tau-V337M organoids. Increased *ELAVL4* expression is therefore consistent with the increased glutamatergic receptor expression and altered splicing we observe in tau-V337M organoids. Dysregulation of *ELAVL4* may not be restricted to FTD-tau; *ELAVL4* is a target of FUS activity, has been found to co-localize with FUS in cytoplasmic speckles and stress granules, and is present in TDP-43 inclusions (Fallini et al., 2012; De Santis et al., 2019). *ELAVL4* dysregulation may therefore be a shared mechanism unifying FTD-tau with other familial forms of FTD.

We demonstrate that susceptibility to glutamate-induced neuronal death in tau-V337M neurons can be blocked by treatment with the PIKFYVE inhibitor, apilimod. It has been proposed that apilimod reduces glutamatergic receptor expression and electrophysiological activity by preventing RAB-dependent recycling of the receptors back to the cell surface. In addition, PIKFYVE inhibition increases the number of EAA1-positive endosomes and LAMP1-positive lysosomes in mouse motor neurons and in human iPSC-derived induced motor neurons (Shi et al., 2018; Staats et al., 2019) and improves proteostasis in neurons *in vivo* (Shi et al., 2018; Staats et al., 2019). Therefore, PIKFYVE inhibition may rescue neurodegeneration by increasing turnover of misfolded tau and favoring elimination of internalized NMDA and AMPA receptors over recycling back to the synapse. Importantly, humans that are haplodeficient for *PIKFYVE* are normal, and apilimod has shown good tolerability in Phase I and II clinical trials, suggesting inhibition could be a viable therapeutic strategy (Sands et al., 2009).

In conclusion, these studies demonstrate the value of applying temporal analysis of human organoid models to the study of neurodegenerative disease to reveal early pathological events. In our working model (Figure 7I), the *MAPT* mutation results in accumulation of mutant tau, which alters synaptic signaling via two mechanisms: altered splicing and homeostasis of glutamatergic signaling genes, and by interfering with proteostasis, which prevents degradation of synaptic glutamatergic receptors. This results in increased glutamatergic receptor expression and maturity, and susceptibility to excitotoxicity. While these changes are initially accommodated by plasticity, over time they result in glutamatergic neuron death. Our discovery that PIKFYVE inhibition, which can dampen glutamate hyperactivity, prevents selective glutamatergic cell death, encourages further study of this pathway as a therapeutic strategy in FTD.

## ACKNOWLEDGMENTS

We are grateful to the Rainwater Charitable Foundation for supporting this collaborative project, to CurePSP for support of organoid protocol development and the New York Genome Center for invaluable help with single cell sequencing. We thank the research subjects and their families who generously participated in this study. We thank Shawn Sutton, Brian Unruh, Isabel Tian, and Nicholas St. John at the NeuraCell core facility https://www.neuracell.org/ for producing organoids, and the Mount Sinai School of Medicine Neuropathology Brain Bank & Research Core for technical assistance. This work was supported by access to equipment made possible by the Hope Center for Neurological Disorders, and the Departments of Neurology and Psychiatry at Washington University School of Medicine. Data collection and dissemination of the data presented in this paper were supported by the LEFFTDS & ARTFL Consortium (LEFFTDS: U01AG045390; ARTFL: U54NS092089). The authors acknowledge the invaluable contributions of the participants in ARTFL & LEFFTDS as well as the assistance of the support staff at each of the participating sites.

## Author Contributions

Conceptualization, K.R.B., M.C.S., J.G., C.M.K, J.K.I., S.J.H., J.F.C., A.M.G., S.T., K.K.

Methodology, K.R.B., M.C.S., J.G., K.S., K.W., S.K.G, S.L., K.L, J.K.I., S.M., J.F.C., C.M.K., D.A.P., N.B

Software, K.R.B., K.W.

Formal Analysis, K.R.B., K.W., T.B., J.G., J.B., J.L., J.K. I., S.M., C.M.K., Y.L., N.B

Investigation, K.R.B., T.B., J.G., K.S., K.W., S.M., J.A.M., C.C., J.B., J.L.

Resources, K.W., S.K.G J.B., J.L., S.L., K.O., C.M.K, S.T.

Data Curation, K.R.B., K.W.

Writing – Original Draft, K.R.B., M.C.S., T.B.

Writing – Review & Editing, K.R.B., M.C.S., B.L.M., C.M.K, J.K.I., S.J.H., J.F.C., A.M.G., S.T., S.M.

Supervision, K.O., B.L.M., C.M.K, J.K.I., S.J.H., J.F.C., A.M.G., S.T.

Project Administration, S.K.G, S.L., K.O., A.M.G., S.T.

Funding Acquisition, K.R.B., M.C.S., K.O., C.M.K, J.K.I., S.J.H., A.M.G., S.T.

## STAR METHODS

### Key Resources Table

| REAGENT or RESOURCE | SOURCE | IDENTIFIER |
| --- | --- | --- |
| **Antibodies** |  |  |
| Total Tau, HT7 (mouse) | Thermo Fisher | MN1000 |
| Total Tau, TAU5 (mouse monoclonal) | Thermo Fisher | AHB0042 |
| Phospho-Tau Ser396 (rabbit polyclonal) | Thermo Fisher | 44752G |
| GAPDH (rabbit monoclonal) | Cell Signaling Technology | 2118S |
| NeuN (mouse monoclonal) | Abcam | 104224 |
| MAP2 (rabbit polyclonal) | Abcam | 32454 |
| LAMP1 (rabbit polyclonal) | Abcam | 24170 |
| GFAP, clone EP672Y (rabbit) | Ventana | 760-4345 |
| Iba1 (rabbit polyclonal) | Invitrogen | PA5-27436 |
| Cathepsin D | Kornfeld lab |  |
| p62 (mouse monoclonal) | Abnova | H00008878-M01 |
| LC3I/II | Novus Biologicals | NB100-2331 |
| PAX6 (rabbit) | BioLegend | 901301 |
| SOX2 (E4) (mouse) | Santa Cruz | sc-365823 |
| FOXG1 (rabbit) | Takara | M227 |
| SOX10 (A-2) (mouse) | Santa Cruz | sc-365692 |
| β-tubulin III (mouse) | Sigma | T-8660 |
| Ki67 (rat IgG2a) | Thermo Fisher | 14-5698-82 |
| Nestin (mouse) | Millipore | MAB5326 |
| MAP2ab (chicken) | Abcam | ab5392 |
| CTIP2 (rat) | Abcam | ab18465 |
| SATB2 (mouse) | Abcam | ab51502 |
| NEUN (mouse) | Millipore | MAB377 |
| GFAP (mouse) | Millipore | MAB3402 |
| S100beta (rabbit) | DAKO Potts | Z0311 |
| Total Tau (rabbit) | DAKO Potts | A-0024 |
| Calbindin 1(rabbit) | Swant | cb-38a |
| VGLUT1 | Gift from Susan Morton, Tom Jessell |  |
| Synaptophysin (mouse) | Abcam | ab8049 |
| PSD95 (rabbit) | Abcam | ab18258 |
| TBR1 (rabbit) | Abcam | ab31940 |
| BRN2 (mouse IgG1) | Millipore | MAB-D51 |
| MEF2C (rabbit) | Novis | NBP2-17260 |
| AT8 (mouse IgG1) | Invitrogen | MN1020 |
| PHF-1 (mouse IgG1) | Gift from Dr. Peter Davies |  |
| Goat anti-rb Alexa 488 | Jackson | 111-545-144 |
| Goat anti-msIgG1 Alexa 546 | Invitrogen | A-21123 |
| Goat anti-mouse IgG2b-Alexa 647 | Invitrogen | A-21242 |
| Goat-mouse IgG1-Alexa 647 | Invitrogen | A-21240 |
| Goat anti-rat IgG2a-Alexa 546 | Invitrogen | A-11081 |
| Goat anti-ck IgG-Alexa 647 | Invitrogen | A-21449 |
| Goat anti-ms IgG1-Alexa647 | Invitrogen | A-21240 |
| **Chemicals, Peptides, and Recombinant Proteins** |  |  |
| Essential 6 medium (E6) | Life Technologies | A1516401 |
| Neurobasal-A | Life Technologies | 10888-022 |
| mTeSR1 medium | StemCell Technologies | 585 |
| Y-27632 | Tocris | 1254 |
| SB431541 | Tocris | 1614 |
| Dorsomorphin | Tocris | 3093 |
| XAV-939 | Tocris | 3748 |
| rhFGF basic | R & D systems | 233-FB |
| human EGF | Peprtech | AF-100-15 |
| human BDNF | Peprtech | 450-02 |
| human NT3 | Peprtech | 450-03 |
| Matrigel | Corning | 356231 |
| B-27 supplement without vitamin A | Life Technologies | 12587010 |
| GlutaMax | Life Technologies | 3505-061 |
| Anti-A | Life Technologies | 15240-062 |
| **Deposited Data** |  |  |
| Bulk RNA-seq data from 2-, 4- and 6-month organoids | GEO |  |
| Single-cell RNA-seq data from 2-, 4- and 6-month organoids | GEO |  |
| **Experimental Models: Cell Lines** |  |  |
| ND32951A.15Δ1B06 (V337V) |  |  |
| ND32951A.15Δ2B09 (V337M) | Tau Consortium | ND-B09 |
| i160311SBB1.1Δ1A02 (V337M) | Tau Consortium | GIH6-A02 |
| i160311SBB1.1Δ1E11 (V337V) | Tau Consortium | GIH6-E11 |
| i160311SBB2.2Δ2A01 (V337M) | Tau Consortium | GIH7-A01 |
| i160311SBB2.2Δ2B12 (V337V) | Tau Consortium | GIH7-B12 |
| i160311SBB2.2Δ2F02 (V337V) | Tau Consortium | GIH7-F02 |
| **Software and Algorithms** |  |  |
| QuPath | Bankhead et al. (2017) QuPath: Open-source software for digital pathology image analysis. Scientific Reports. https://doi.org/10.1038/s41598-017-17204-5 | https://qupath.github.io/ |
| Seurat 3.0 (dev) | Stuart, T. et al (2019). Comprehensive integration of single cell data. Cell. https://doi.org/10.1016/j.cell.2019.05.031 | https://satijalab.org/seurat/ |
|  | Butler, A. et al. (2018). Integrating single-cell transcriptomic data across different conditions, technologies, and species. Nature Biotechnology. https://doi.org/10.1038/nbt.4096 |  |
| Monocle3 | Trapnell, C. et al. (2014). The dynamics and regulators of cell fate decisions are revealed by pseudotemporal ordering of single cells. Nature Biotechnology. https://doi.org/10.1038/nbt.2859<br>Qiu, X. et al. (2017). Reversed graph embedding resolves complex single-cell trajectories. Nature Methods. https://doi.org/10.1038/nmeth.4402 | https://cole-trapnell-lab.github.io/monocle3/ |
| SingleR | Aran, D. et al (2019). Reference-based analysis of lung single-cell sequencing reveals a transitional profibrotic macrophage. Nature Immunology. https://doi.org/10.1038/s41590-018-0276-y | https://bioconductor.org/packages/devel/bioc/vignettes/SingleR/inst/doc/SingleR.html |
| Script for differential gene expression over pseudotime trajectories | Kanton, S. et al (2019). Organoid single-cell genomic atlas uncovers human-specific features of brain development. Nature. https://doi.org/10.1038/s41586-019-1654-9 |  |
| **Other** |  |  |
| GW17-18 cortical scRNA-seq data | Polioudakis, D. et al (2019). A Single-Cell Transcriptomic Atlas of Human Neocortical Development during Mid-gestation. Neuron. https://doi.org/10.1016/j.neuron.2019.06.011 | http://solo.bmap.ucla.edu/shiny/webapp/ |
| Adult astrocyte bulk RNAseq data | Zhang, Y. et al (2016). Purification and Characterization of Progenitor and Mature Human Astrocytes Reveals Transcriptional and Functional Differences with Mouse. Neuron. https://doi.org/10.1016/j.neuron.2015.11.013 | https://www.brainrnaseq.org/ |
| ELAVL RIP-seq data | Scheckel (2016). Regulatory consequences of neuronal ELAV-like protein binding to coding and non-coding RNAs in human brain eLife. https://doi.org/10.7554/eLife.10421 |  |

### RESOURCE AVAILABILITY

#### Lead Contacts

Further information and requests for resources and reagents should be directed to and will be fulfilled by the Lead Contacts, Sally Temple and Alison Goate.

#### Materials Availability

Human iPSC lines used in this study are available for request from the Tau Consortium via the Neural Stem Cell Institute (https://www.neuralsci.org/tau).

#### Data and Code Availability

The bulk and single cell RNA-sequencing data generated as part of this study are available at the Gene Expression Omnibus (GEO; https://www.ncbi.nlm.nih.gov/geo/). Previously unpublished code for ELAVL4 binding cargo enrichment can be found in the supplemental information. All other code used in this study have been previously published, and the pipelines described in the Method Details.

### EXPERIMENTAL MODEL AND SUBJECT DETAILS

#### Cell lines

Human iPSC lines obtained from the Tau Consortium cell line collection (https://www.neuralsci.org/tau) (Karch et al., 2019) GIH6-1-C1ΔE11 (WT/WT), GIH6-1-C1ΔA02 (V337M/WT), GIH7-C2Δ2B12 (WT/WT), GIH7-C2Δ2F02 (WT/WT), GIH7-C2Δ2A01 (V337/WT), ND32951A.15Δ1B06 (WT/WT), and ND32951A.15Δ1B09 (V337M/WT), NeuraCell, NY, USA; Table S1 (all female lines)) were maintained in mTeSR1 medium based on feeder-free culture protocols in six-well plates (Corning) coated with growth factor-reduced Matrigel. To ensure consistency and obtain high quality organoid production, feeder free (FF) maintained cultures were grown to a density not higher than 80% with daily feeding. Passaging of iPSCs colonies was carried out using Dispase to the desired split ratio, and cells were incubated at 37°C and 5% CO2. Cells were maintained with daily medium changes of 2 ml mTeSR per well.

### METHOD DETAILS

#### Preparation of iPSCs for 3D organoid production

Organoids were generated as previously described (Yoon et al., 2019). Human iPSCs were grown to 80~85% confluency prior to AggreWell^™^800 (StemCell Tech. cat no #34815) induction. E8 medium was supplemented with 10μM ROCK inhibitor - Y-27632 to make EB formation medium. EB formation medium was added to each well of the AggreWell plate and centrifuged at 2,000 x g for 5 minutes in a swinging bucket rotor fitted with a plate holder to remove any air bubbles from the microwells. The microwells were checked under a microscope to ensure all bubbles had been removed. A single cell suspension of iPSCs was prepared by pre-treating the iPSCs for 50-60 minutes with EB formation medium. The maintenance medium was aspirated from the iPSC culture plates and rinsed twice with DPBS (Dulbecco’s Phosphate Buffered Solution) (without Calcium or Magnesium). 2 ml of Accutase was added per well of a 6 well plate and incubated for ~ 10 minutes at 37°C in 5% CO_2_ until cells detached from the dish by gentle shaking. The cells were counted and centrifuged at 1,200 rpm for 4 minutes. 3 million cells were transferred into individual wells of the AggreWell^™^800 plate, to achieve 10,000 cells per microwell. Gentle pipetting of the cell suspension in the microwells was done to ensure an even distribution of cells throughout the well. The AggreWell^™^800 plate was centrifuged at 100xg for 3 minutes to distribute the cells into the microwell cones and checked under a microscope to verify even distribution of cells among the microwell cones. The cells were incubated at 37°C and 5% CO_2_ for 24 hours.

#### 3D organoid patterning and growth

DMEM/F12 and Essential 6 (E6) media, were pre-warmed to room temperature. HiPSC-derived spheroids were removed from the microwells by firmly pipetting medium in the wells up and down 2-3 times with a P1000 pipette and transferred to a 50 ml conical tube and washed with DMEM/F12. The washed organoids were transferred to ultra-low attachment 10 cm plates (Corning, catalog #3262) in 10mL E6 medium supplemented with 2.5 μM Dorsomorphin (DM), 10 μM SB431542, and 2.5 μM XAV-939. Media changes were performed daily for five days. On the sixth day in suspension, the spheroids were transferred to neural medium (NM), 1x B-27 supplement without vitamin A, 1x GlutaMax, 1x Anti-A, 20 ng/ml FGF2 and 20 ng/ml EGF for 19 days, with daily medium changes for the first 10 days, and every other day for the subsequent 9 days. To promote differentiation of neural progenitors into neurons, FGF2 and EGF were replaced with 20 ng/ml BDNF and 20 ng/ml NT3 (from day 25). From day 43 onwards, medium changes were done every four days using only NM without growth factors.

#### Fixation and Frozen Sectioning of Organoids

Organoids were fixed overnight at 4°C in 4% paraformaldehyde, rinsed three times in PBS and placed in 30% sucrose in PBS for three days. Organoids were then placed in cryomolds (Tissue Tek Catalog #4565) with OCT compound (Tissue Tek, catalog # 4583), snap frozen and stored at minus 80°C. Organoids were cryostat sectioned sequentially at 20μm thickness using a Leica cryostat (model CM3050S). Sections were placed on microscope glass slides, dried overnight and stored at −20°C for subsequent immunohistochemistry.

#### Immunohistochemistry of Organoid Frozen Sections

Slides were thawed to room temperature, and lines drawn to partition the edges of sections on the glass slides using a PAPpen (ImmEdge Pen; Vector Labs, Catalog #H4000). The sections were rehydrated, blocked, permeabilized and immunostained with primary antibodies: PAX6 (1:200), SOX2 (1:100), β-tubulin III (1:1000), FOXG1 (1:500), SOX10 (1:100), Ki67 (1:200), Nestin (1:200), GFAP (1:400), S100beta (1:400), CTIP2 (1:500), SATB2 (1:50), MAP2AB (1:1000), NeuN (1:100), TBR1(1:500), BRN2 (1:500), MEF2C (1:200), AT8(1:1000), PHF1(1:1000), Total Tau (1:200), VGLUT1(1:16,000), Calbindin1 (1:10,000), PSD95 (1:1000) and Synaptophysin (1:200). Primary antibodies were incubated overnight at 4°C, washed three times with PBS and then incubated with corresponding Alexa Fluor conjugated secondary antibody (1:1000) for 1 hour at room temperature. Antibody manufacturer information is shown in Key Resources Table. Sections were coverslipped and imaged using fluorescence and confocal microscopy (Zeiss AXIO Observer.Z1; Zeiss 780).

#### Immunofluorescence Image Analysis of Frozen Sections

Fluorescence image analysis on immunostained frozen sections was performed using a custom macro developed for ImageJ software. Low-power images were used to visualize across organoid sections. Individual thresholds were set to measure CTIP2 and DAPI cell counts within each image. CTIP2/DAPI values were calculated and normalized to the 2 month samples within an individual organoid batch and cell line. Box plots generated contain counts from at least three organoids per batch and across three different organoid batches.

#### Immunohistochemistry of Organoid Paraffin Sections

Organoids were fixed in 10% neutral buffered formalin and embedded in paraffin blocks at the Neuropathology Brain Bank CoRE (NPBB) at the Icahn School of Medicine at Mount Sinai. Sections (4-5 μm) were prepared from organoid blocks, mounted on positively charged slides and baked overnight at 70°C. IHC was performed on a Benchmark XT autostainer platform (Ventana) according to the manufacturer’s protocol with reagents and antibodies acquired from the same lot. Antigen retrieval was done using CC1 (citric acid buffer, Roche Diagnostics, Basel Switzerland) for 1 hour. All primary antibodies were diluted in Antibody dilution buffer (ABD24, Roche Diagnostics). Antibody manufacturer information shown in Key Resources Table. Primary antibodies were incubated for either 16 minutes (GFAP 1:10), 24 minutes (MAP2 1:50), 32 minutes (ALDH1L1 1:100), 40 minutes (Iba1 1:100) or 44 minutes (NeuN 1:100) at 37°C. Primary antibodies were visualized using either the Ultraview or Optiview detection kit (Roche Diagnostics) as an indirect biotin-free system to detect primary antibodies according to manufacturer’s directions. All slides were counterstained with hematoxylin for 8 minutes and coverslipped. Slides were visualized using a Nikon Eclipse brightfield microscope with a Nikon DS-Fi3 camera and NIS-Elements software. For every set of slides stained, a neuropathologically confirmed severe AD case and healthy control case was included as a batch control.

#### Immunohistochemistry Semi-quantitative and Digital quantitative morphometric analysis

Pathological changes assessed using semi-quantitative analysis were scored by two researchers (JFC and KW) blinded to mutational and control conditions using a semi-quantitative scoring system as follows: no staining (0), low staining (1), moderate staining (2) or high staining (3). For unbiased digital quantitative assessment, slides were imaged using an Aperio CS2 (Leica Biosystems, Wetzlar Germany) digital slide scanner at the Department of Pathology at Mount Sinai. Individual organoids were manually segmented into individual annotations on whole slide images (WSI) sections, batch-processed and analyzed in QuPath (version 0.2.3, https://QuPath.github.io/). All analysis was batch-processed using a custom analysis workflow and cell-based analysis was applied to individual organoids using the following methodology. Neuronal density was measured by the number of NeuN+ cells detected using a positive cell detection classifier based on thresholded values for DAB intensity and detected object area. All quantitative values were normalized to area.

#### Bulk RNA sequencing

##### Organoid preparation and RNA extraction

Individual organoids were collected and stored in RNAlater (Qiagen) at −80°C from each time point across 3 batches of differentiation. RNA was then extracted in a single batch using the KingFisher Flex (ThermoFisher Scientific) automated system and MagMAX mirVana Total RNA isolation kit (ThermoFisher Scientific). The resulting RNA was submitted to GeneWiz for 150bp unstranded paired-end poly-A RNA sequencing on the Illumina platform.

##### Data alignment and analysis

Paired-end RNA sequencing reads were trimmed for Illumina TruSeq adapters and aligned to the human hg38 genome using the STAR aligner (Dobin et al., 2013) for quantification of gene-level read counts (FastQC v0.11.8 used for quality control of reads, Cutadapt v2.5 for adapter trimming, STAR v2.5.2b for read alignment). Genes were filtered by expression abundance (counts per million, CPM, > 0.1 in 5 or more samples), and library sizes were estimated by the trimmed mean of M-values normalization (TMM) method (Liao et al., 2014) with the R edgeR package v3.26.8. Differential gene expression between *MAPT* mutation organoids and isogenic controls was predicted by linear mixed model analysis using the variancePartition package (v1.14.1) in the R programming environment (Hoffman and Roussos, 2020) to account for sample correlations within isogenic lines and differentiation batches. The false discovery rate (FDR) of the differential expression test was estimated using the Benjamini-Hochberg (BH) method (Reiner et al., 2003). Functional Gene Set Enrichment Analysis (GSEA) was performed using the Molecular Signatures Database (MSigDB) Gene Ontology (GO) annotations of canonical pathway, biological process, cellular compartment and molecular function, with 10,000 permutations to establish FDR values (implemented with the R GOtest package v1.0.8 (https://rdrr.io/github/mw201608/GOtest/)). Additional pathway analyses were carried out using Ingenuity Pathway Analysis (QIAGEN Inc., https://www.qiagenbioinformatics.com/products/ingenuitypathway-analysis). LeafCutter v0.2.7 (Li et al., 2018) was used to identify differentially spliced intron clusters between Tau-V337M and Tau-V337V organoids at each time point using default settings. Analysis of GO enrichment terms for differentially spliced genes was carried out using g:Profiler (https://biit.cs.ut.ee/gprofiler/gost).

#### Cell Hashing Single-Cell RNA-seq

##### Organoid dissociation and Cell Hashing

Organoids were incubated in Papain (Worthington) for 90-120 minutes at 37°C with gentle shaking (300rpm) and regular pipetting with a glass Pasteur pipette to obtain single cell suspensions. Cells were pelleted and resuspended in ice-cold PBS before being passed through a 40μm filter and kept on ice. Viability and cell number was determined by Trypan Blue staining and counting using the Countess system (ThermoFisher Scientific). 50-100,000 cells per organoid were submitted to the New York Genome Center (NYGC) for hashtag-oligonucleotide (HTO) antibody binding, library preparation and sequencing as previously described (Stoeckius et al., 2017, 2018).

##### Single-Cell RNA-seq data QC and alignment

Data QC and alignment was carried out at the NYGC. Briefly, analysis up to the counting of UMIs per cell per gene was carried out using Drop-Seq tools (Drop-seq tools v1.0, McCarroll Lab), and reads were aligned to the human hg38 genome using the STAR aligner (Dobin et al., 2013). HTOs were identified and counted using Cite-Seq Count (https://hoohm.github.io/CITE-seq-Count/), and the samples were demultiplexed using HTODemux (Butler et al., 2018). Following processing at NYGC, the data was further processed using Seurat 3.0 dev (Butler et al., 2018; Stuart et al., 2019) to filter out cells with <200 detectable genes, <500 reads and >20% mitochondrial rate. We retrieved 8000-10,000 high quality cells per lane with a read depth of ~25,000 reads per cell.

##### Data analysis

Cells were annotated using the SingleR package (Aran et al., 2019), with custom reference libraries constructed from human fetal neocortical single-cell RNA sequencing data (Polioudakis et al., 2019) and bulk adult astrocyte RNA-sequencing data (Zhang et al., 2016). The top 3000 most variable genes at each time point were used as input for cell type annotation. Data from different sequencing runs, batches and ages were integrated and scaled using SCTransform (Hafemeister and Satija, 2019), while regressing out the following covariates; percentage of mitochondrial genes, the number of cells collected per parent organoid, the number of genes per cell and the number of reads per cell. Data was also corrected for differentiation batch. Using Seurat, principal components analysis (PCA) was carried out using the top 3,000 most variable genes, and data reduction was performed with UMAP (McInnes et al., 2018). For glutamatergic neuron and astrocyte analyses, individual cell types were isolated from each cell line and integrated across each time point using the same covariates as described above. Data from each cell type was then re-scaled and reduced for each isogenic pair. Clusters were identified using the default resolution factor, and mutant-cell enriched clusters were identified as those with a log2FC > 0.5 greater proportion of V337M cells compared to V337V cells. Differential gene expression analysis was carried out on raw count data between mutant-enriched clusters and remaining clusters using the MAST model with appropriate covariates. Significantly differentially expressed genes were submitted to pathway analysis in IPA, and pathways shared between all isogenic pairs and altered in the same direction were identified. Cells in V337M and V337V organoids were separately ordered in pseudotime using Monocle3 (Qiu et al., 2017; Trapnell et al., 2014) using the top 3000 most variable genes. Differential expression between V337M and V337V glutamatergic neurons was carried out using a custom analysis as described previously (Kanton et al., 2019), using a spline function with 6 degrees of freedom. Genes were clustered based on trajectory using the R package pheatmap (Kolde, 2019) using the ward.D2 method, and genes with differential expression trajectories were submitted for GO enrichment using g:Profiler (https://biit.cs.ut.ee/gprofiler/gost).

#### Intersection of ELAVL4-bound genes

We obtained a list of genes previously demonstrated to be bound by ELAVL in human brain using RNA immunoprecipitation RNA-seq (Scheckel et al., 2016). The ELAVL4-RIP genes were intersected with the genes we previously identified to be differentially spliced. Using the ‘hypeR’ package, we then explored the functions of the bound genes by calculating the GOBP enriched categories followed by semantic analysis with the ‘rrvgo’ package. Code for this analysis can be found in the Supplemental Information.

#### Western blotting analysis of tau and P-tau

Organoids were removed from the cell culture media, placed in microcentrifuge tubes and rapidly frozen on dry ice. The organoids were lysed using RIPA buffer (50 mM Tris, 150mM sodium chloride, 0.5% sodium deoxycholate, 2% SDS, 1% nonidet P-40, Boston Bioproducts, Ashland, MA) containing 1× concentration of HALT protease and phosphatase inhibitors (Thermo Fisher Scientific, Waltham, MA). Briefly, the RIPA buffer was added to the tubes containing organoids and incubated on ice for 20 min followed by manual disruption and lysis using a wide bore pipette tip. Next, the lysate was sonicated in water at 4°C for 20 min (20 sec on, 20 sec off) followed by centrifugation at 14,000 × g at 4°C. The supernatant was collected and protein concentration was determined using the BCA assay (Thermo Fisher). A total of 5 μg protein was separated by SDS-PAGE (7% Tris-acetate gel) according to standard methods. Proteins were transferred from the gel onto PVDF membranes (EMD Millipore) using standard procedures. Membranes were blocked in 5% (w/v) BSA (Sigma) in Tris-buffered saline with 0.1% Tween-20 (TBST, Boston Bio-Products), incubated overnight with primary antibody (Key Resources Table) at 4°C, followed by corresponding HRP-linked secondary antibody at 1:4000 dilution (Cell Signaling Technology). Blots were developed with SuperSignal Chemiluminescent Substrate (ThermoFisher) according to manufacturer’s instructions, exposed to autoradiographic film (LabScientific by ThermoFisher), and scanned on an Epson Perfection V800 Photo Scanner. Protein bands’ densitometry (pixel mean intensity in arbitrary units, a.u.) was measured with Image J and normalized to the respective internal control (GAPDH) band.

#### Immunoblotting for autophagy-lysosomal markers

Organoids were extracted in lysis buffer (50mM Tris pH 7.6, 1mM EDTA, 150 mM NaCl, 1% TritonX-100, phosphatase and protease inhibitors (Millipore-Sigma)) and incubated on ice for 5 minutes. Lysates were then centrifuged at 14,000xg for 10 minutes at 4°C, and the resulting supernatant was saved for analysis. Total protein levels were assayed by bicinchoninic acid assay (BCA) assay (Thermo-Fisher). Standard sodium dodecyl sulfate-polyacrylamide gel electrophoresis (SDS-PAGE) was performed in 4-12% Criterion Tris-HCl gels (Bio-Rad) with 10 μg of total protein loaded in each well. Samples were boiled in Laemmli sample buffer prior to electrophoresis. Gels were transferred to PVDF and the resulting immunoblots were probed with the antibodies listed above. Antibodies were visualized with SuperSignal West Pico Chemiluminescent Substrate (Thermo) or Lumigen ECL Ultra (TMA-6) according to manufacturer’s instructions. Immunoblots were exposed on a Syngene G:Box iChemi XT Gel Documentation System and imaged using GeneSnap software according to manufacturer’s instructions. Band intensity was analyzed using GeneTools, normalized against total protein by Ponceau S staining of each blot, and band intensity was expressed relative to the normalized control within each blot.

#### Longitudinal Tracking of Neuron Survival in 3-D

Cerebral organoids were transduced with lentivirus (with 1:1000 polybrene) encoding *Synapsin1*::eGFP for 5 days and fixed in Matrigel (VWR) prior to experimental use. Glutamate (5 mM) was administered to the media on day zero during the survival time course, and replenished after one, two, four, and six days. Continuous z-stacks spanning 150 μM from the organoid surface were captured daily using a Zeiss AxioZoom.v16 wide-field upright fluorescent microscope. Post-processing and image alignment were performed using ImageJ software (NIH).

### QUANTIFICATION AND STATISTICAL ANALYSIS

Statistical analysis for tau expression and phosphorylation were performed using two-tailed unpaired student’s t-test. \**p* < 0.05, \*\**p* <0.01, \*\*\**p* <0.001, *n* = 9 per group from three independent experiments. All tests were performed using GraphPad Prism V 8. Statistical significance was determined at 0.05. For lysosomal assays, protein quantification from three technical replicates and 3 biological replicates were averaged and expressed as mean ± standard error of the mean (SEM). Statistical difference was measured using an unpaired Student’s t-test. For quantification of lysosomal protein analyte levels in the tau-V337M organoids and isogenic controls, graph represents mean ± SEM. Significance was determined using an unpaired, t-test. \**p* < 0.05. Statistical analysis and data visualization was performed using GraphPad Prism v9.0.0 (La Jolla, CA).

Statistical analysis of CTIP2-positive neurons via immunofluorescence was performed using 2-way ANOVA. \**p* < 0.05, \*\*\*\**p* < 0.0001. Organoids were analyzed across 3 unique organoid batches, number of organoids analyzed per sample are reported in figure (*n* = number of organoids/biological replicates). Quantifiable phenotypes by immunohistochemistry were compared between mutants and controls at 2, 4 and 6 months using a Mann-Whitney U test (Figure 2C, NeuN, 4 months: n_mutant_ = 24, n_isogenic_ = 25, Mann-Whitney U = 175, p = 0.0119, 6 months: n_mutant_ = 26, n_isogenic_ = 34, Mann-Whitney U = 274, p = 0.0117; Figure S2B, MAP2, 6 months, n_mutant_ = 29, n_isogenic_ = 33, Mann-Whitney U = 303.5, p = 0.0060). All statistical analysis and data visualization was performed using GraphPad Prism v8.4.3 (La Jolla, CA). Statistical analysis on survival tracking of individual neurons (n) in mutant and corrected isogenic control organoids in Figures 7B (n_isogenic_= 80, n_mutant_= 80, Chi-sq = 3.723, *p* = 0.0537), 7D (n_isogenic_= 80, n_mutant_= 80, Chi-sq = 17.77, *p* < 0.0001) 7G (n_mutant+DMSO_= 80, n_mutant+apilimod_= 80, Chi-sq = 17.25, *p* < 0.0001), and 7H (n_isogenic+DMSO_= 80, n_mutant+apilimod_= 80, Chi-sq =.3797, *p* = 0.3797) were performed using Log-rank (Mantel-Cox) tests. Significance defined as *p* < 0.05. Statistical analysis and data visualization was performed using GraphPad Prism v9.0.0 (La Jolla, CA).

The analysis pipelines of bulk and single-cell RNA sequencing data are described in the Method Details. 241 individual organoids were collected and submitted for bulk RNA-sequencing across all cell lines, three time points and three differentiation batches. Read quality was assessed using MultiQC **(https://multiqc.info/).** Two organoids were excluded from further analysis due to inconsistent positioning away from the remaining samples by principal components analysis, resulting in a final dataset consisting of 239 organoids (V337M *n* = 99, V337V *n* = 140). All presented differential expression, GSEA, pathway enrichment and splicing analyses p-values were adjusted for multiple test correction. For single cell RNA-sequencing, 348 individual organoids were processed (V337M *n* = 151, V337V *n* = 197), of which 339 passed quality control, determined by samples containing detectable cells with clear HTO identities. Additional filtering is described in Method Details. The resulting dataset for analysis included 379,064 cells (V337M *n* = 161,706, V337V *n* = 217,358). All reported p-values were adjusted for multiple test correction. Significance was defined as adjusted *p* < 0.05.

## SUPPLEMENTAL INFORMATION

**Table S1.** Human iPSC lines employed to generate 3D cerebral organoids.

| MAPT | Age at Biopsy | iPSC ID | Organoid ID | Abbreviated ID | Sanger seq. confirmed. | Karyotype |
| --- | --- | --- | --- | --- | --- | --- |
| V337M | 40s | ND32951A.15Δ2B09 | ND32951A.15Δ1B09 | NDB09 | yes | normal |
| V337V | 40s | ND32951A.15Δ1B06 | ND32951A.15Δ1B06 | NDB06 | yes | normal |
| V337M | 60s | i160311SBB1.1Δ1A02 | GIH-6-C1-ΔA02 | GIH6-A02 | yes | normal |
| V337V | 60s | i160311SBB1.1Δ1E11 | GIH-6-C1-Δ1E11 | GIH6-E11 | yes | normal |
| V337M | 30s | i160311SBB2.2Δ2A01 | GIH-7-C2-Δ2A01 | GIH7-A01 | yes | normal |
| V337V | 30s | i160311SBB2.2Δ2B12 | GIH-7-C2-Δ2B12 | GIH7-B12 | yes | normal |
| V337V | 30s | i160311SBB2.2Δ2F02 | GIH-7-C2-Δ2F02 | GIH7-F02 | yes | normal |

### Supplementary Figures Legends

**Figure S1.**
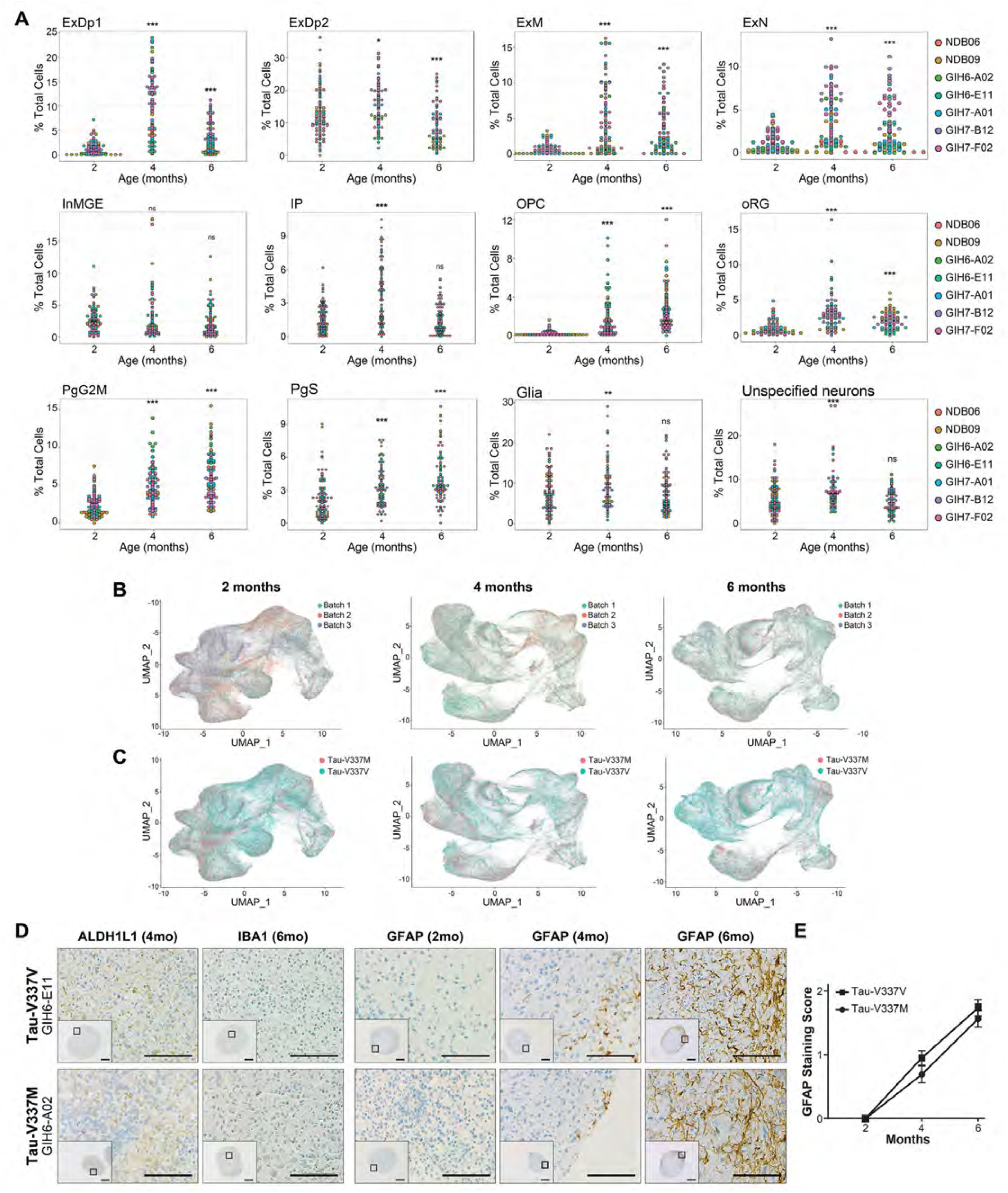
Forebrain Organoid Differentiation Recapitulates Key Features of Human Brain Patterning, Related to Figure 1. **(A)** Cell type proportions and variability for individual organoids over time for all cell types identified. Points are colored by donor line. Differences in cell type proportion are calculated compared to proportion of cells at 2 months, *p<0.05, **p<0.01, ***p<0.001. **(B-C)** UMAP reduction of cell hashing single cell sequencing data from forebrain organoids at 2, 4 and 6 months of age, colored by batch **(B)** and mutation **(C)**. **(D)** Immunohistochemical staining of glial markers ALDH1L1 (4-month organoids), IBA1 (6-month) and GFAP (2, 4, and 6 months) in tau-V337M (GIH6-A02) and isogenic corrected (GIH6-E11) organoids. Scale bars 250 μm (inserts) and 50 μm. **(E)** Quantification of GFAP+ neurons at 2, 4 and 6 months of organoid differentiation in tau-V337M (GIH6-A02) and isogenic corrected (GIH6-E11) organoids.

**Figure S2.**
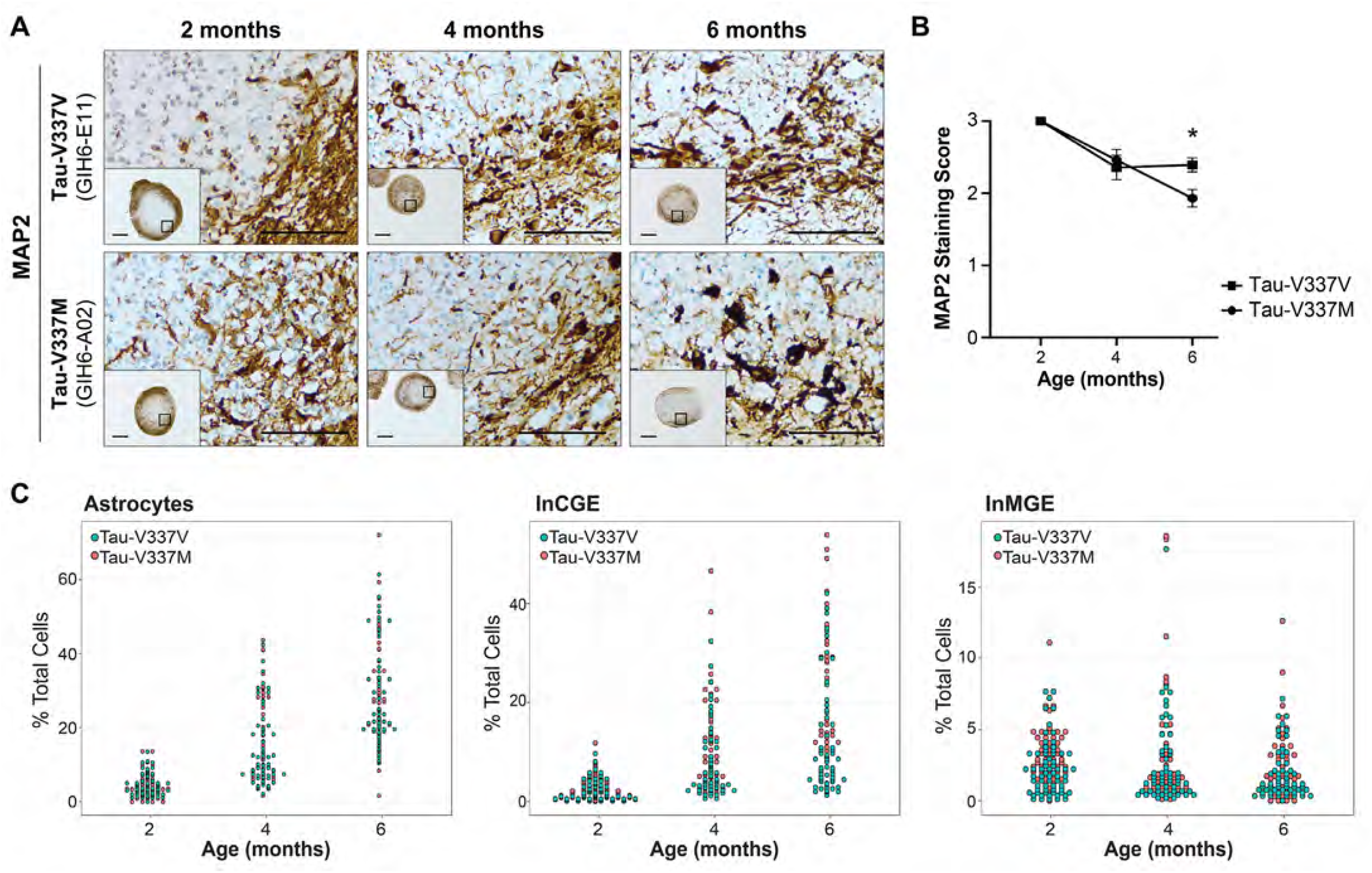
Tau-V337M Organoids Exhibit Neuronal-Specific Loss Over Time, Related to Figure 2. **(A-B)** Imaging of MAP2+ neurons and quantitative analysis at 2, 4 and 6 months, in mutant (GIH6-A02) and isogenic corrected (GIH6-E11) organoids. Statistical analysis by Mann-Whitney: at 6 months \*\**p* = 0.006. **(C)** Proportion of Astrocytes, InCGE (inhibitory neurons from the caudal ganglionic eminence), and InMGE (inhibitory neurons from the medial ganglionic eminence) per organoid at 2, 4 and 6 months of differentiation. Points are colored by mutation (V337M = red, V337V = blue). A linear model was conducted between glutamatergic cell type proportions in mutant vs. isogenic-corrected organoids at each time point, *p* > 0.05, not significant.

**Figure S3.**
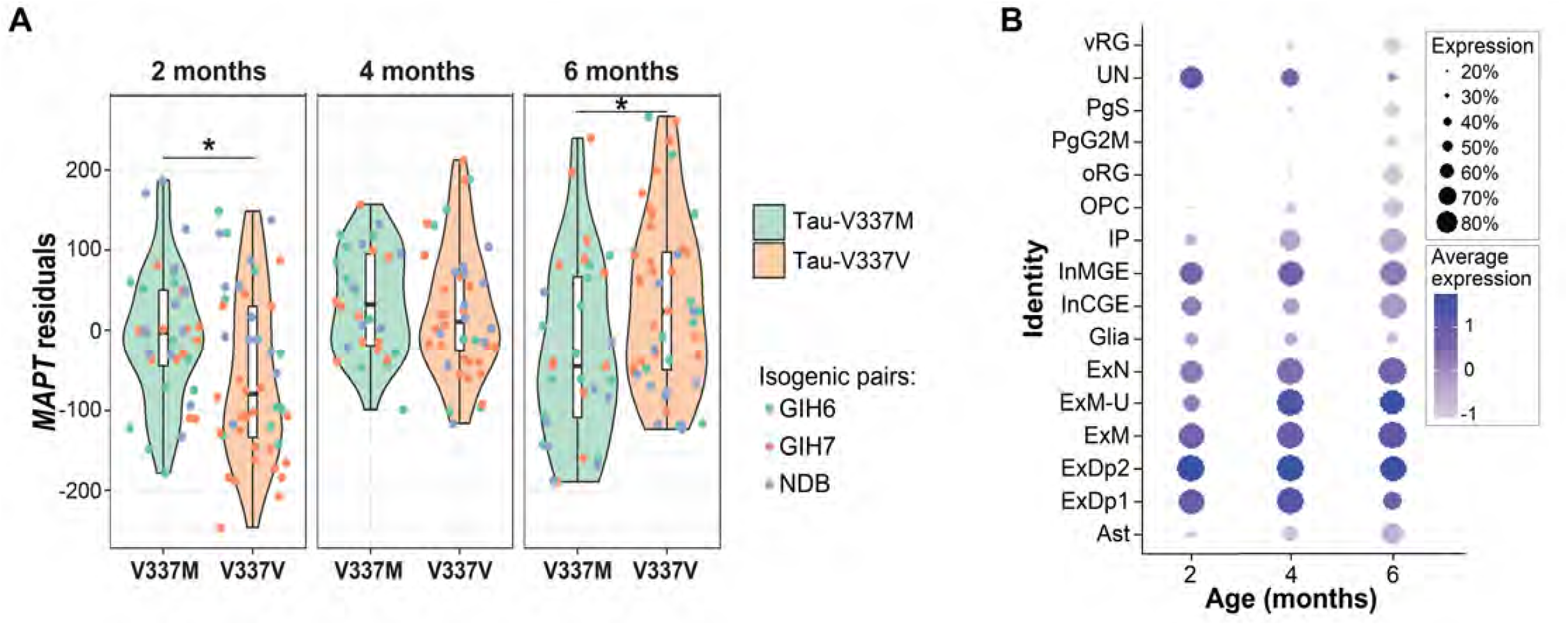
Characterization of *MAPT* Expression in Organoids, Related to Figure 4. **(A)** Violin plots show residuals of *MAPT* expression in organoids derived from bulk RNA-seq data following correction for covariates (V337M = green, V337V = orange). Each isogenic cell line is denoted by different color data points (GIH6 = green, GIH7 = orange, NDB = purple). Statistical comparisons (linear mixed model for repeated measures) were carried out between V337M and V337V organoids at each time point, \**p* < 0.05. **(B)** Proportion of *MAPT*-expressing cell-types in 2, 4 and 6-month organoids, with expression scaled within each time point. Dot size represents the proportion of cells expressing *MAPT*, and depth of color denotes level of *MAPT* expression.

**Figure S4.**
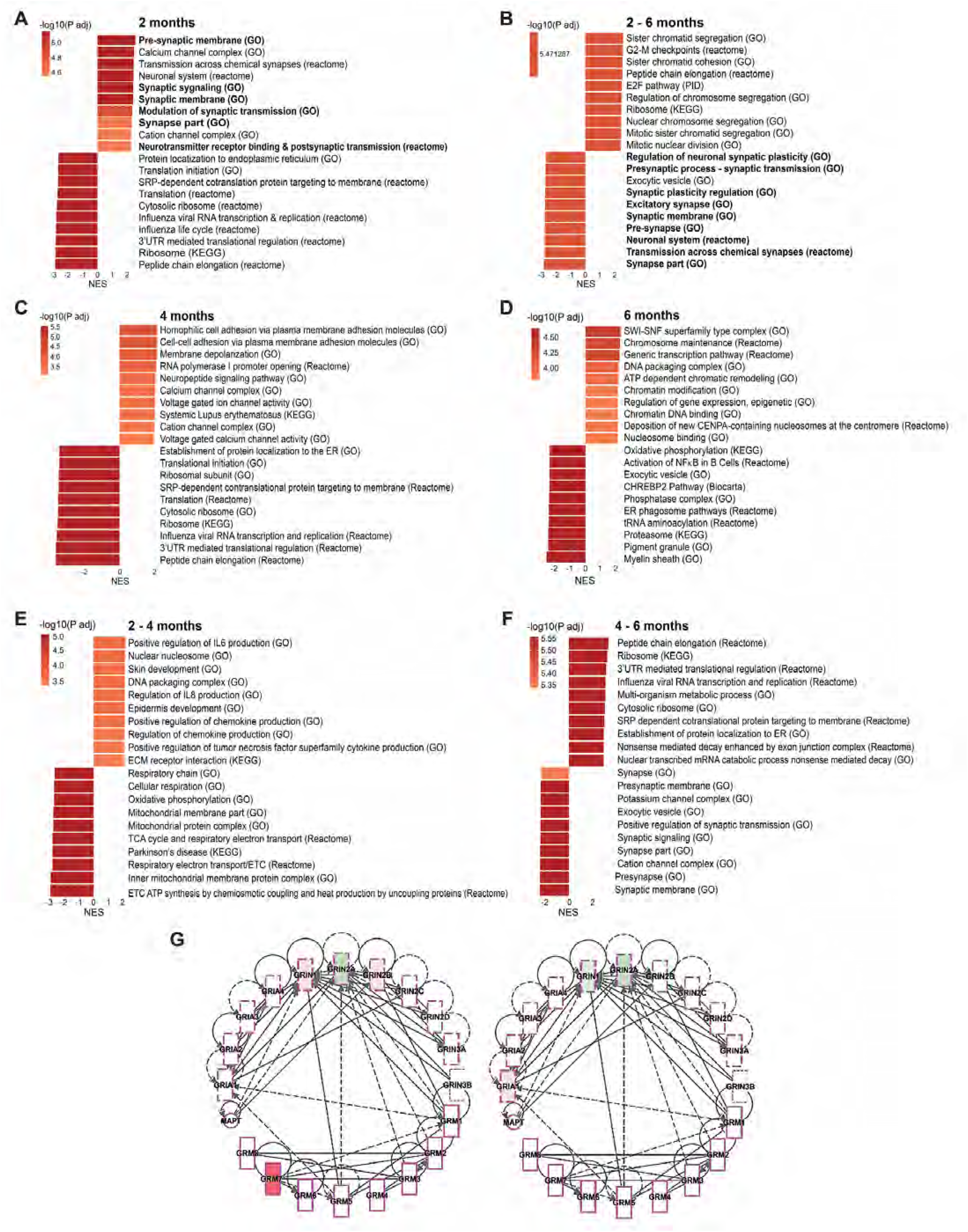
Changes in Enriched Gene Expression Pathways in Tau-V337M Organoids, Related to Figure 5. **(A-B)** Gene set enrichment analysis of differentially expressed genes in the bulk RNA-seq data at 2 months of differentiation **(A)** and the interaction between mutation and age from 2-6 months **(B).** NES = normalized enrichment score. Adjusted-log10 *p*-value is indicated by depth of color on bars. Synaptic-related pathways are highlighted in bold. **(C-F)** Gene set enrichment analysis of differentially expressed genes in the bulk RNA-seq data at 4 months **(C),** 6 months **(D)** and the interaction between mutation and age between 2-4 months **(E)** and 4-6 months **(F).** NES = normalized enrichment score. Adjusted-log10 *p*-value is indicated by depth of color on bars. **(G)** Expression and connectivity of glutamatergic receptor genes and *MAPT* at 4 months (left) and 6 months (right) of organoid differentiation. Red indicates gene upregulation in V337M organoids compared to isogenic controls, and green indicates gene downregulation. Depth of color reflects the extent of fold-change expression.

**Figure S5.**
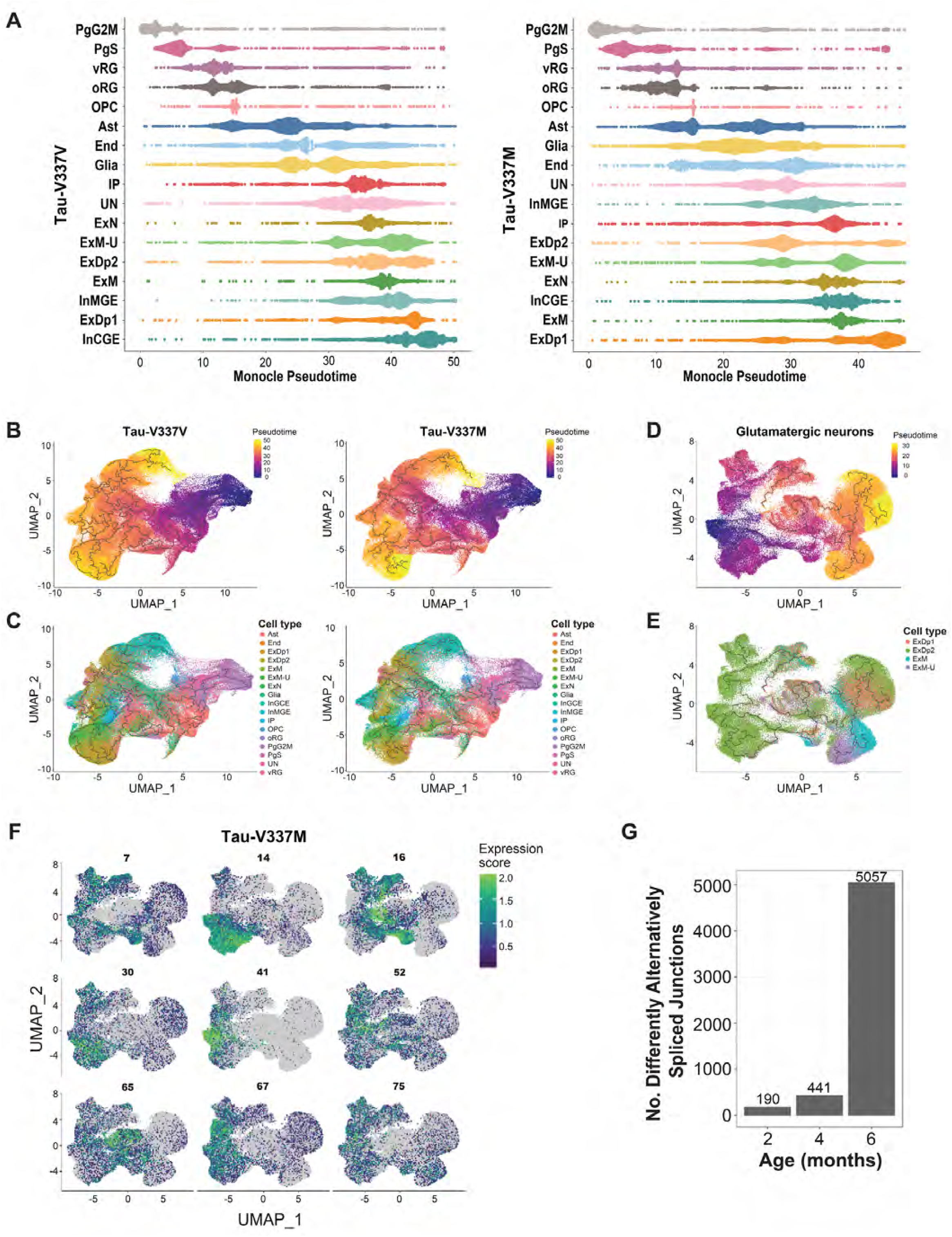
Ordering of Organoid Cell Types and Enrichment of Tau-V337M and V337V Gene Expression Modules in Pseudotime, Related to Figure 6. **(A)** Separation of cell types present in organoids by pseudotime in tau-V337V (left) and tau-V337M (right) organoids. **(B-C)** UMAP reduction of all V337V (left) and V337M (right) cells colored by pseudotime **(B)** and cell type **(C).** Cells appearing earliest in pseudotime are denoted by dark purple, and those latest in pseudotime are in yellow. **(D-E)** UMAP reduction of all excitatory neurons colored by pseudotime **(D)** and cell type **(E).** Cells earliest in pseudotime are denoted by dark purple, and those latest in pseudotime are in yellow. **(F)** UMAPs of excitatory neurons indicating the location and expression score of the top 9 V337M-enriched gene clusters. **(G)** Number of significantly differentially spliced intron junctions between tau-V337M and tau-V337V organoids at each differentiation time-point (exact number shown above each bar).

## REFERENCES

Akamatsu, W., Fujihara, H., Mitsuhashi, T., Yano, M., Shibata, S., Hayakawa, Y., Okano, H.J., Sakakibara, S.I., Takano, H., Takano, T., et al. (2005). The RNA-binding protein HuD regulates neuronal cell identity and maturation. Proc. Natl. Acad. Sci. U. S. A. 102, 4625–4630.

Apicco, D.J., Zhang, C., Maziuk, B., Jiamg, L., Ballance, H., Boudeau, S., Ung, C., Li, H., and Wolozin, B. (2019). Dysregulation of RNA Splicing in Tauopathies Daniel. Cell Rep. 29, 4377–4388.

Aran, D., Looney, A., Liu, L., Wu, E., Fong, V., Hsu, A., Chak, A., Naikawadi, R., Wolters, P., Abate, A., et al. (2019). Reference-based analysis of lung single-cell sequencing reveals a transitional profibrotic macrophage. Nat. Immunol. 20, 163–172.

Bain, H.D.C., Davidson, Y.S., Robinson, A.C., Ryan, S., Rollinson, S., Richardson, A., Jones, M., Snowden, J.S., Pickering-Brown, S., and Mann, D.M.A. (2019). The role of lysosomes and autophagosomes in frontotemporal lobar degeneration. Neuropathol. Appl. Neurobiol. 45, 244–261.

Bankhead, P., Loughrey, M.B., Fernández, J.A., Dombrowski, Y., McArt, D.G., Dunne, P.D., McQuaid, S., Gray, R.T., Murray, L.J., Coleman, H.G., et al. (2017). QuPath: Open source software for digital pathology image analysis. Sci. Rep. 7, 1–7.

Beagle, A., Darwish, S., Ranasinghe, K., La, A., Karageorgiou, E., and Vossel, K. (2017). Relative Incidence of Seizures and Myoclonus in Alzheimer’s Disease, Dementia with Lewy Bodies, and Frontotemporal Dementia Alexander. J. Alzheimer’s Dis. 60, 221–223.

Bendiske, J., and Bahr, B.A. (2003). Lysosomal activation is a compensatory response against protein accumulation and associated synaptopathogenesis - An approach for slowing Alzheimer disease? J. Neuropathol. Exp. Neurol. 62, 451–463.

Benussi, A., Padovani, A., and Borroni, B. (2015). Phenotypic heterogeneity of monogenic frontotemporal dementia. Front. Aging Neurosci. 7, 1–19.

Benussi, A., Alberici, A., Buratti, E., Ghidoni, R., Gardoni, F., Luca, M. Di, Padovani, A., and Borroni, B. (2019). Toward a glutamate hypothesis of frontotemporal dementia. Front. Neurosci. 13, 1–9.

Bhaduri, A., Andrews, M.G., Mancia, W., Jung, D., Allen, D., Jung, D., Schmunk, G., Haeussler, M., Pollen, A.A., Nowakowski, T.J., et al. (2020). Cell Stress in Cortical Organoids Impairs Molecular Subtype Specification. Nature 578, 142–148.

Bird, T., Knopman, D., VanSwieten, J., Rosso, S., Feldman, H., Tanabe, H., Graff-Raford, N., Geschwind, D., Verpillat, P., and Hutton, M. (2003). Epidemiology and genetics of frontotemporal dementia/Pick’s Disease. Ann. Neurol. 54, 29–31.

Bodea, L.G., Eckert, A., Ittner, L.M., Piguet, O., and Götz, J. (2016). Tau physiology and pathomechanisms in frontotemporal lobar degeneration. J. Neurochem. 138, 71–94.

Bolognani, F., Contente-Cuomo, T., and Perrone-Bizzozero, N.I. (2009). Novel recognition motifs and biological functions of the RNA-binding protein HuD revealed by genome-wide identification of its targets. Nucleic Acids Res. 38, 117–130.

Borroni, B., Stanic, J., Verpelli, C., Mellone, M., Bonomi, E., Alberici, A., Bernasconi, P., Culotta, L., Zianni, E., Archetti, S., et al. (2017). Anti-AMPA GluA3 antibodies in Frontotemporal dementia: A new molecular target. Sci. Rep. 7, 1–10.

Borroni, B., Benussi, A., Premi, E., Alberici, A., Marcello, E., Gardoni, F., Di Luca, M., and Padovani, A. (2018). Biological, Neuroimaging, and Neurophysiological Markers in Frontotemporal Dementia: Three Faces of the Same Coin. J. Alzheimer’s Dis. 62, 1113–1123.

Bowie, D. (2008). Ionotropic Glutamate Receptors & CNS Disorders. CNS Neurol. Disord. - Drug Targets 7, 129–143.

Bronicki, L.M., and Jasmin, B.J. (2013). Emerging complexity of the HuD/ELAVl4 gene; Implications for neuronal development, function, and dysfunction. Rna 19, 1019–1037.

Butler, A., Hoffman, P., Smibert, P., Papalexi, E., and Satija, R. (2018). Integrating single-cell transcriptomic data across different conditions, technologies, and species. Nat. Biotechnol. 36.

Cai, X., Xu, Y., Cheung, A.K., Tomlinson, R.C., Alcázar-, A., Murphy, L., Billich, A., Zhang, B., Feng, Y., Klumpp, M., et al. (2016). PIKfyve, a class III PI-kinase, is the target of the small molecular IL12/23 inhibitor apilimod and a new player in toll-like receptor signaling. 20, 912–921.

Camp, J.G., Badsha, F., Florio, M., Kanton, S., Gerber, T., Wilsch-Bräuninger, M., Lewitus, E., Sykes, A., Hevers, W., Lancaster, M., et al. (2015). Human cerebral organoids recapitulate gene expression programs of fetal neocortex development. Proc. Natl. Acad. Sci. U. S. A. 112, 15672–15677.

Clare, R., King, V., Wirenfeldt, M., and Vinters, H. (2010). Synapse Loss in Dementias. J Neurosci Res 88, 2083–2090.

DeBoer, E.M., Azevedo, R., Vega, T.A., Brodkin, J., Akamatsu, W., Okano, H., Wagner, G.C., and Rasin, M.R. (2014). Prenatal deletion of the RNA-binding protein HuD disrupts postnatal cortical circuit maturation and behavior. J. Neurosci. 34, 3674–3686.

Dobin, A., Davis, C.A., Schlesinger, F., Drenkow, J., Zaleski, C., Jha, S., Batut, P., Chaisson, M., and Gingeras, T.R. (2013). STAR: Ultrafast universal RNA-seq aligner. Bioinformatics 29, 15–21.

Ehrlich, M., Hallmann, A.L., Reinhardt, P., Araúzo-Bravo, M.J., Korr, S., Röpke, A., Psathaki, O.E., Ehling, P., Meuth, S.G., Oblak, A.L., et al. (2015). Distinct Neurodegenerative Changes in an Induced Pluripotent Stem Cell Model of Frontotemporal Dementia Linked to Mutant TAU Protein. Stem Cell Reports 5, 83–96.

Fallini, C., Bassell, G.J., and Rossoll, W. (2012). The ALS disease protein TDP-43 is actively transported in motor neuron axons and regulates axon outgrowth. Hum. Mol. Genet. 21, 3703–3718.

Faravelli, I., Costamagna, G., Tamanini, S., and Corti, S. (2020). Back to the origins: Human brain organoids to investigate neurodegeneration. Brain Res. 1727, 146561.

Fong, H., Wang, C., Knoferle, J., Walker, D., Balestra, M.E., Tong, L.M., Leung, L., Ring, K.L., Seeley, W.W., Karydas, A., et al. (2013). Genetic correction of tauopathy phenotypes in neurons derived from human induced pluripotent stem cells. Stem Cell Reports 1, 226–234.

Forrest, S.L., Kril, J.J., Stevens, C.H., Kwok, J.B., Hallupp, M., Kim, W.S., Huang, Y., McGinley, C. V., Werka, H., Kiernan, M.C., et al. (2018). Retiring the term FTDP-17 as MAPT mutations are genetic forms of sporadic frontotemporal tauopathies. Brain 141, 521–534.

Ghetti, B., Oblak, A.L., Boeve, B.F., Johnson, K.A., Dickerson, B.C., and Goedert, M. (2015). Invited review: Frontotemporal dementia caused by microtubule-associated protein tau gene (MAPT) mutations: A chameleon for neuropathology and neuroimaging. Neuropathol. Appl. Neurobiol. 41, 24–46.

Gonzalez, C., Armijo, E., Bravo-Alegria, J., Becerra-Calixto, A., Mays, C.E., and Soto, C. (2018). Modeling amyloid beta and tau pathology in human cerebral organoids. Mol. Psychiatry 23, 2363–2374.

Hafemeister, C., and Satija, R. (2019). Normalization and variance stabilization of single-cell RNA-seq data using regularized negative binomial regression. Genome Biol. 20, 1–15.

Hoffman, G.E., and Roussos, P. (2020). Dream: powerful differential expression analysis for repeated measures designs. Bioinformatics.

Ince-dunn, G., Okano, H.J., Jensen, K., Park, W., Ule, J., Mele, A., Fak, J., Yang, C., Zhang, C., Yoo, J., et al. (2012). Neuronal Elav-like (Hu) proteins regulate RNA splicing and abundance to control glutamate levels and neuronal excitability. 75, 1067–1080.

Iovino, M., Agathou, S., Gonz??lez-Rueda, A., Del Castillo Velasco-Herrera, M., Borroni, B., Alberici, A., Lynch, T., O’Dowd, S., Geti, I., Gaffney, D., et al. (2015). Early maturation and distinct tau pathology in induced pluripotent stem cell-derived neurons from patients with MAPT mutations. Brain 138, 3345–3359.

Jiang, S., Wen, N., Li, Z., Dube, U., Del Aguila, J., Budde, J., Martinez, R., Hsu, S., Fernandez, M. V., Cairns, N.J., et al. (2018). Integrative system biology analyses of CRISPR-edited iPSC-derived neurons and human brains reveal deficiencies of presynaptic signaling in FTLD and PSP. Transl. Psychiatry 8, 1–16.

Kanton, S., Boyle, M.J., He, Z., Santel, M., Weigert, A., Sanchís-calleja, F., Guijarro, P., Sidow, L., Fleck, J.S., Han, D., et al. (2019). Human-Specific Features of Brain Development.

Karch, C.M., Kao, A.W., Karydas, A., Onanuga, K., Martinez, R., Argouarch, A., Wang, C., Huang, C., Sohn, P.D., Bowles, K.R., et al. (2019). A Comprehensive Resource for Induced Pluripotent Stem Cells from Patients with Primary Tauopathies. Stem Cell Reports 13.

Koike, M., Tsukada, S., Tsuzuki, K., Kijima, H., and Ozawa, S. (2000). Regulation of kinetic properties of GluR2 AMPA receptor channels by alternative splicing. J. Neurosci. 20, 2166–2174.

Kolde, R. (2019). pheatmap: Pretty Heatmaps. R package version 1.0.12.

Lancaster, M.A., Renner, M., Martin, C.A., Wenzel, D., Bicknell, L.S., Hurles, M.E., Homfray, T., Penninger, J.M., Jackson, A.P., and Knoblich, J.A. (2013). Cerebral organoids model human brain development and microcephaly. Nature 501, 373–379.

Lancaster, M.A., Corsini, N.S., Wolfinger, S., Gustafson, E.H., Phillips, A.W., Burkard, T.R., Otani, T., Livesey, F.J., and Knoblich, J.A. (2017). Guided self-organization and cortical plate formation in human brain organoids. Nat. Biotechnol. 35, 659–666.

Leuzy, A., Zimmer, E.R., Dubois, J., Pruessner, J., Cooperman, C., Soucy, J.P., Kostikov, A., Schirmaccher, E., Désautels, R., Gauthier, S., et al. (2016). In vivo characterization of metabotropic glutamate receptor type 5 abnormalities in behavioral variant FTD. Brain Struct. Funct. 221, 1387–1402.

Li, Y., Knowles, D., Humphrey, J., Barbeira, A., Dickinson, S., Im, H., and Pritchard, J. (2018). Annotation-free quantification of RNA splicing using LeafCutter. Nat. Genet. 50, 151–158.

Liao, Y., Smyth, G.K., and Shi, W. (2014). FeatureCounts: An efficient general purpose program for assigning sequence reads to genomic features. Bioinformatics 30, 923–930.

Lin, L.C., Nana, A.L., Hepker, M., Hwang, J.H.L., Gaus, S.E., Spina, S., Cosme, C.G., Gan, L., Grinberg, L.T., Geschwind, D.H., et al. (2019). Preferential tau aggregation in von Economo neurons and fork cells in frontotemporal lobar degeneration with specific MAPT variants. Acta Neuropathol. Commun. 7, 1–10.

Lin, W.-L., Lewis, J., Yen, S.-H., Hutton, M., and Dickson, D.W. (2003). Ultrastructural neuronal pathology in transgenic mice expressing mutant (P301L) human tau. J Neurocytol 32.

Liu, H., Wang, H., Peterson, M., Zhang, W., Hou, G., and Zhang, Z. wei (2019). N-terminal alternative splicing of GluN1 regulates the maturation of excitatory synapses and seizure susceptibility. Proc. Natl. Acad. Sci. U. S. A. 116, 21207–21212.

Marat, A.L., and Haucke, V. (2016). Phosphatidylinositol 3-phosphates—at the interface between cell signalling and membrane traffic. EMBO J. 35, 561–579.

McInnes, L., Healy, J., and Melville, J. (2018). UMAP: Uniform manifold approximation and projection for dimension reduction. ArXiv.

Nakamura, M., Shiozawa, S., Tsuboi, D., Amano, M., Watanabe, H., Maeda, S., Kimura, T., Yoshimatsu, S., Kisa, F., Karch, C.M., et al. (2019). Pathological Progression Induced by the Frontotemporal Dementia-Associated R406W Tau Mutation in Patient-Derived iPSCs. Stem Cell Reports 13, 684–699.

Nixon, R.A., Wegiel, J., Kumar, A., Yu, W.H., Peterhoff, C., Cataldo, A., and Cuervo, A.M. (2005). Extensive involvement of autophagy in Alzheimer disease: An immuno-electron microscopy study. J. Neuropathol. Exp. Neurol. 64, 113–122.

Olszewska, D.A., Lonergan, R., Fallon, E.M., and Lynch, T. (2016). Genetics of Frontotemporal Dementia. Curr. Neurol. Neurosci. Rep. 16.

Paşca, A.M., Sloan, S.A., Clarke, L.E., Tian, Y., Makinson, C.D., Huber, N., Kim, C.H., Park, J.-Y., O’Rourke, N.A., Nguyen, K.D., et al. (2015). Functional cortical neurons and astrocytes from human pluripotent stem cells in 3D culture. Nat. Methods 12, 671–678.

Pei, W., Huang, Z., Wang, C., Han, Y., Jae, S.P., and Niu, L. (2009). Flip and flop: A molecular determinant for AMPA receptor channel opening. Biochemistry 48, 3767–3777.

Piras, A., Collin, L., Grüninger, F., Graff, C., and Rönnbäck, A. (2016). Autophagic and lysosomal defects in human tauopathies: analysis of post-mortem brain from patients with familial Alzheimer disease, corticobasal degeneration and progressive supranuclear palsy. Acta Neuropathol. Commun. 4, 22.

Polioudakis, D., de la Torre-Ubieta, L., Langerman, J., Elkins, A.G., Shi, X., Stein, J.L., Vuong, C.K., Nichterwitz, S., Gevorgian, M., Opland, C.K., et al. (2019). A Single-Cell Transcriptomic Atlas of Human Neocortical Development during Mid-gestation. Neuron 103, 785–801.e8.

Qiu, X., Mao, Q., Tang, Y., Wang, L., Chawla, R., Pliner, H., and Trapnell, C. (2017). Reversed graph embedding resolves complex single-cell trajectories. Nat. Methods 14, 979–982.

Ravanidis, S., Kattan, F.G., and Doxakis, E. (2018). Unraveling the pathways to neuronal homeostasis and disease: Mechanistic insights into the role of RNA-binding proteins and associated factors. Int. J. Mol. Sci. 19, 1–49.

Reiner, A., Yekutieli, D., and Benjamini, Y. (2003). Identifying differentially expressed genes using false discovery rate controlling procedures. Bioinformatics 19, 368–375.

Rohrer, J.D., Guerreiro, R., Vandrovcova, J., Uphill, J., Reiman, D., Beck, J., Isaacs, A.M., Authier, A., Ferrari, R., Fox, N.C., et al. (2009). The heritability and genetics of frontotemporal lobar degeneration. Neurology 73, 1451–1456.

Sands, B.E., Jacobson, E.W., Sylwestrowicz, T., Younes, Z., Dryden, G., Fedorak, R., and Greenbloom, S. (2009). Randomized, double-blind, placebo-controlled trial of the oral interleukin-12/23 inhibitor apilimod mesylate for treatment of active Crohn’s disease. Inflamm. Bowel Dis. 16, 1209–1218.

De Santis, R., Alfano, V., de Turris, V., Colantoni, A., Santini, L., Garone, M.G., Antonacci, G., Peruzzi, G., Sudria-Lopez, E., Wyler, E., et al. (2019). Mutant FUS and ELAVL4 (HuD) Aberrant Crosstalk in Amyotrophic Lateral Sclerosis. Cell Rep. 27, 3818–3831.e5.

Sato, C., Barthélemy, N.R., Mawuenyega, K.G., Patterson, B.W., Gordon, B.A., Jockel-Balsarotti, J., Sullivan, M., Crisp, M.J., Kasten, T., Kirmess, K.M., et al. (2018). Tau Kinetics in Neurons and the Human Central Nervous System. Neuron 97, 1284–1298.e7.

Scheckel, C., Drapeau, E., Frias, M.A., Park, C.Y., Fak, J., Zucker-Scharff, I., Kou, Y., Haroutunian, V., Ma’ayan, A., Buxbaum, J.D., et al. (2016). Regulatory consequences of neuronal ELAV-like protein binding to coding and non-coding RNAs in human brain. Elife 5, 1–35.

Seebohm, G., Neumann, S., Theiss, C., Novkovic, T., Hill, E. V., Tavaré, J.M., Lang, F., Hollmann, M., Manahan-Vaughan, D., and Strutz-Seebohm, N. (2012). Identification of a novel signaling pathway and its relevance for GluA1 recycling. PLoS One 7.

Shi, Y., Lin, S., Staats, K., Li, Y., Chang, W.-H., Hung, S.-T., Hendricks, E., Linares, G., Wang, Y., Son, E., et al. (2018). Haploinsufficiency Leads to Neurodegeneration in C9ORF72 ALS/FTD Human Induced Motor Neurons. Nat. Med. 24, 313–325.

Shi, Y., Hung, S.T., Rocha, G., Lin, S., Linares, G.R., Staats, K.A., Seah, C., Wang, Y., Chickering, M., Lai, J., et al. (2019). Identification and therapeutic rescue of autophagosome and glutamate receptor defects in C9ORF72 and sporadic ALS neurons. JCI Insight 4, 1–21.

Shiarli, A.M., Jennings, R., Shi, J., Bailey, K., Davidson, Y., Tian, J., Bigio, E.H., Ghetti, B., Murrell, J.R., Delisle, M.B., et al. (2006). Comparison of extent of tau pathology in patients with frontotemporal dementia with Parkinsonism linked to chromosome 17 (FTDP-17), frontotemporal lobar degeneration with Pick bodies and early onset Alzheimer’s disease. Neuropathol. Appl. Neurobiol. 32, 374–387.

Silva, M.C., Cheng, C., Mair, W., Almeida, S., Fong, H., Biswas, M.H.U., Zhang, Z., Huang, Y., Temple, S., Coppola, G., et al. (2016). Human iPSC-Derived Neuronal Model of Tau-A152T Frontotemporal Dementia Reveals Tau-Mediated Mechanisms of Neuronal Vulnerability. Stem Cell Reports 7, 325–340.

Silva, M.C., Nandi, G.A., Tentarelli, S., Gurrell, I.K., Jamier, T., Lucente, D., Dickerson, B.C., Brown, D.G., Brandon, N.J., and Haggarty, S.J. (2020). Prolonged tau clearance and stress vulnerability rescue by pharmacological activation of autophagy in tauopathy neurons. Nat. Commun. 11.

Sohn, P.D., Huang, C.T.L., Yan, R., Fan, L., Tracy, T.E., Camargo, C.M., Montgomery, K.M., Arhar, T., Mok, S.A., Freilich, R., et al. (2019). Pathogenic Tau Impairs Axon Initial Segment Plasticity and Excitability Homeostasis. Neuron 104, 458–470.e5.

Spillantini, M.G., Crowther, R., and Goedert, M. (1996). Comparison of the neurofibrillary pathology in Alzheimer’s disease and familial presenile dementia with tangles. Acta Neuropathol. 92, 42–48.

Spina, S., Schonhaut, D.R., Boeve, B.F., Seeley, W.W., Ossenkoppele, R., Neil, J.P.O., Lazaris, A., Rosen, H.J., Boxer, A.L., Perry, D.C., et al. (2017). Frontotemporal dementia with the V337M MAPT mutation Tau-PET and pathology correlations. Neurology 1–10.

Sposito, T., Preza, E., Mahoney, C.J., Set??-Salvia, N., Ryan, N.S., Morris, H.R., Arber, C., Devine, M.J., Houlden, H., Warner, T.T., et al. (2015). Developmental regulation of tau splicing is disrupted in stem cell-derived neurons from frontotemporal dementia patients with the 10 + 16 splice-site mutation in MAPT. Hum. Mol. Genet. 24, 5260–5269.

Staats, K.A., Seah, C., Sahimi, A., Wang, Y., Koutsodendris, N., Lin, S., Kim, D., Chang, W.H., Gray, K.A., Shi, Y., et al. (2019). Small molecule inhibition of PIKFYVE kinase rescues gain- and loss-of-function C9ORF72 ALS/FTD disease processes in vivo. BioRxiv.

Stoeckius, M., Hafemeister, C., Stephenson, W., Houck-loomis, B., Chattopadhyay, P.K., Swerdlow, H., Satija, R., and Smibert, P. (2017). Simultaneous epitope and transcriptome measurement in single cells. Nat. Methods 14.

Stoeckius, M., Zheng, S., Houck-loomis, B., Hao, S., Yeung, B.Z., Iii, W.M.M., Smibert, P., and Satija, R. (2018). Cell Hashing with barcoded antibodies enables multiplexing and doublet detection for single cell genomics. Genome Biol. 1–12.

Stuart, T., Butler, A., Hoffman, P., Hafemeister, C., Papalexi, E., Mauck, W.M., Hao, Y., Stoeckius, M., Smibert, P., and Satija, R. (2019). Comprehensive Integration of Single-Cell Data. Cell 177, 1888–1902.e21.

Sumi, S.M., Bird, T.D., Nochlin, D., and Raskind, M.A. (1992). Familial presenile dementia with psychosis associated with cortical neurofibrillary tangles and degeneration of the amygdala. Neurology 42, 120–127.

Tebaldi, T., Zuccotti, P., Peroni, D., Köhn, M., Gasperini, L., Potrich, V., Bonazza, V., Dudnakova, T., Rossi, A., Sanguinetti, G., et al. (2018). HuD Is a Neural Translation Enhancer Acting on mTORC1-Responsive Genes and Counteracted by the Y3 Small Non-coding RNA. Mol. Cell 71, 256–270.e10.

Tiruchinapalli, D.M., Caron, M.G., and Keene, J.D. (2008). Activity-dependent expression of ELAV/Hu RBPs and neuronal mRNAs in seizure and cocaine brain. J. Neurochem. 107, 1529–1543.

Trapnell, C., Cacchiarelli, D., Grimsby, J., Pokharel, P., Morse, M., Lennon, N.J., Livak, K.J., Mikkelsen, T.S., Rinn, L., and Biology, R. (2014). Pseudo-temporal ordering of individual cells reveals dynamics and regulators of cell fate decisions Cole. 32, 381–386.

Trevino, A.E., Sinnott-Armstrong, N., Andersen, J., Yoon, S.J., Huber, N., Pritchard, J.K., Chang, H.Y., Greenleaf, W.J., and Pașca, S.P. (2020). Chromatin accessibility dynamics in a model of human forebrain development. Science (80-.). 367.

Vanderweyde, T., Apicco, D.J., Youmans-Kidder, K., Ash, P., Cook, C., Lummertz da Rocha, E., Jansen-West, K., Frame, A., Citro, A., Leszyk, J., et al. (2016). Interaction of tau with the RNA-Binding Protein TIA1 Regulates tau Pathophysiology and Toxicity. Cell Rep. 15, 1455–1466.

Velasco, S., Kedaigle, A.J., Simmons, S.K., Nash, A., Rocha, M., Quadrato, G., Paulsen, B., Nguyen, L., Adiconis, X., Regev, A., et al. (2019). Individual brain organoids reproducibly form cell diversity of the human cerebral cortex. Nature 570, 523–527.

Verheyen, A., Diels, A., Reumers, J., Van Hoorde, K., Van den Wyngaert, I., van Outryve d’Ydewalle, C., De Bondt, A., Kuijlaars, J., De Muynck, L., De Hoogt, R., et al. (2018). Genetically Engineered iPSC-Derived FTDP-17 MAPT Neurons Display Mutation-Specific Neurodegenerative and Neurodevelopmental Phenotypes. Stem Cell Reports 11, 363–379.

Wang, Y., Martinez-Vicente, M., Krüger, U., Kaushik, S., Wong, E., Mandelkow, E.M., Cuervo, A.M., and Mandelkow, E. (2009). Tau fragmentation, aggregation and clearance: The dual role of lysosomal processing. Hum. Mol. Genet. 18, 4153–4170.

Warmus, B.A., Sekar, D.R., McCutchen, E., Schellenberg, G.D., Roberts, R.C., McMahon, L.L., and Roberson, E.D. (2014). Tau-mediated NMDA receptor impairment underlies dysfunction of a selectively vulnerable network in a mouse model of frontotemporal dementia. J. Neurosci. 34, 16482–16495.

Watanabe, M., Buth, J.E., Vishlaghi, N., Taxidis, J., Khakh, B., Coppola, G., Pearson, C.A., Gong, D., Dai, X., Damoiseaux, R., et al. (2017). Self-organized cerebral organoids with human specific features predict effective drugs to combat Zika virus infection. 21, 517–532.

Weyn-Vanhentenryck, S.M., Feng, H., Ustianenko, D., Duffié, R., Yan, Q., Jacko, M., Martinez, J.C., Goodwin, M., Zhang, X., Hengst, U., et al. (2018). Precise temporal regulation of alternative splicing during neural development. Nat. Commun. 9.

Wray, S. (2017). Modeling tau pathology in human stem cell derived neurons. Brain Pathol. 27, 525–529.

Wren, M.C., Zhao, J., Liu, C.C., Murray, M.E., Atagi, Y., Davis, M.D., Fu, Y., Okano, H.J., Ogaki, K., Strongosky, A.J., et al. (2015). Frontotemporal dementia-associated N279K tau mutant disrupts subcellular vesicle trafficking and induces cellular stress in iPSC-derived neural stem cells. Mol. Neurodegener. 10, 1–13.

Yoon, S.-J., Elahi, L., Pasca, A., Marton, R., Gordon, A., O, R., Miura, Y., Walczak, E., Holdgate, G., Fan, H., et al. (2019). Reliability of human 3D cortical organoid generation. Nat. Methods 16, 75–78.

Yu, W.H., Kumar, A., Peterhoff, C., Shapiro Kulnane, L., Uchiyama, Y., Lamb, B.T., Cuervo, A.M., and Nixon, R.A. (2004). Autophagic vacuoles are enriched in amyloid precursor protein-secretase activities: Implications for β-amyloid peptide over-production and localization in Alzheimer’s disease. Int. J. Biochem. Cell Biol. 36, 2531–2540.

Zhang, Y., Sloan, S.A., Clarke, L.E., Caneda, C., Plaza, C.A., Blumenthal, P.D., Vogel, H., Steinberg, G.K., Edwards, M.S.B., Iii, J.A.D., et al. (2016). Purification and characterization of progenitor and mature human astrocytes reveals transcriptional and functional differences with mouse. Neuron 89, 37–53.

Zhou, H.L., Hinman, M.N., Barron, V.A., Geng, C., Zhou, G., Luo, G., Siegel, R.E., and Lou, H. (2011). Hu proteins regulate alternative splicing by inducing localized histone hyperacetylation in an RNA-dependent manner. Proc. Natl. Acad. Sci. U. S. A. 108, 627–635.

Zhu, H., Hinman, M.N., Hasman, R.A., Mehta, P., and Lou, H. (2008). Regulation of Neuron-Specific Alternative Splicing of Neurofibromatosis Type 1 Pre-mRNA. Mol. Cell. Biol. 28, 1240–1251.

